# Diaminopurine in Nonenzymatic RNA Template Copying

**DOI:** 10.1101/2024.02.20.581310

**Authors:** Xiwen Jia, Ziyuan Fang, Seohyun Chris Kim, Dian Ding, Lijun Zhou, Jack W. Szostak

## Abstract

In the primordial RNA World, before the advent of ribozymes, nonenzymatic template copying would have been essential for the transmission of genetic information. However, the products of chemical copying with the canonical nucleotides are heavily biased towards the incorporation of G and C. Diaminopurine (D) can form a D:U base pair that is almost as strong as a G:C base pair. We therefore asked whether replacing A with D might lead to more efficient and less biased nonenzymatic template copying. As expected, primer extension substrates containing D bind to U in the template more tightly than substrates containing A. However, primer extension with D exhibited elevated reaction rates on a C template, leading to concerns about fidelity. To investigate the nature of the D:C mismatch, we solved the crystal structure of RNA duplexes containing D:C mismatches, and showed that D can form a wobble-type base pair with C. We then asked whether competition with G would decrease mismatched primer extension. We performed nonenzymatic primer extension with all four activated nucleotides on randomized RNA templates containing all four letters, and used deep sequencing to analyze the products. We found that the DUCG genetic system exhibited a more even product distribution and a lower mismatch frequency than the canonical AUCG system. Furthermore, primer extension is greatly reduced following all mismatches, including the D:C mismatch. Our study suggests that diaminopurine deserves further attention for its possible role in the RNA World, and as a potentially useful component of artificial nonenzymatic RNA replication systems.

## Introduction

The RNA World hypothesis posits RNA as the primordial genetic polymer due to its dual role in encoding information and catalyzing reactions.^1–3^ Prior to the emergence of macromolecular catalysts such as ribozymes, nonenzymatic template copying likely played a critical role in the transmission of hereditary information.^4^ This process, however, has a marked tendency to favor the incorporation of guanosine (G) and cytidine (C) nucleotides over adenosine (A) and uridine (U),^5^ presenting a bias in the copying process. To address this imbalance, we have looked beyond the four canonical nucleotides found in RNA, seeking alternatives that could mitigate this issue.

Diaminopurine (D), an adenine analogue characterized by an additional exocyclic amine, is a potentially prebiotic nucleobase. It has been detected in carbonaceous meteorites^6^, albeit at low parts per billion (ppb) levels. Subsequent studies have demonstrated the synthesis of the D deoxynucleoside (β-2,6-diaminopurine 2′-deoxyriboside)^7^ by transglycoslyation of 2-thiouridine with diaminopurine, although with a low yield of 2%. A more efficient synthesis of 2,6-diaminopurine ribonucleoside 2′-phosphate from ribose 1′-2′ cyclic phosphate and 2,6-diaminopurine was demonstrated by Kim and Benner^8^, suggesting that D nucleotides could have formed on the early Earth if a high yielding synthesis of these precursors was possible. Moreover, D and its derivatives, including deoxyribonucleosides and DNA trimers containing D, have a photostability that is only two-fold less than that of A, implying that D nucleotides could have withstood the UV radiation flux at the surface of the early Earth.^7,9^ Collectively, these findings support the further study of potentially prebiotic pathways for the origin and accumulation of D nucleotides, as well as motivating the continued study of potential roles for D in the formation and replication of prebiotic RNA or DNA oligonucleotides.

In addition to its potential prebiotic relevance, diaminopurine exhibits molecular properties that may have provided early evolutionary advantages through the stabilization of nucleic acid duplexes. In RNA duplexes, D forms three hydrogen bonds when paired with U, as opposed to two in the A:U pair (Figure 1).^10,11^ This, together with a stronger stacking interaction, results in a D:U base pair being energetically more favorable than an A:U base pair.^12^ Consequently, there is an increased free energy change upon duplex formation, with a net ΔΔG°_37_ of -0.29 kcal/mol per A:U to D:U base pair substitution, as predicted by a combined molecular dynamics/quantum mechanics (MD/QM) approach.^12^ The larger degree of stabilization is also evidenced by a significant rise in the experimentally determined melting temperature (T_m_) of D:U RNA duplexes compared to A:U duplexes.^13^ This enhanced duplex stability extends to other sugar backbones such as DNA^14,15^ and threose nucleic acid (TNA)^16^. However, within the context of nonenzymatic RNA replication, the increased duplex stability from the stronger D:U pair may introduce challenges by increasing the difficulty of strand separation.^17^

**Figure 1.**
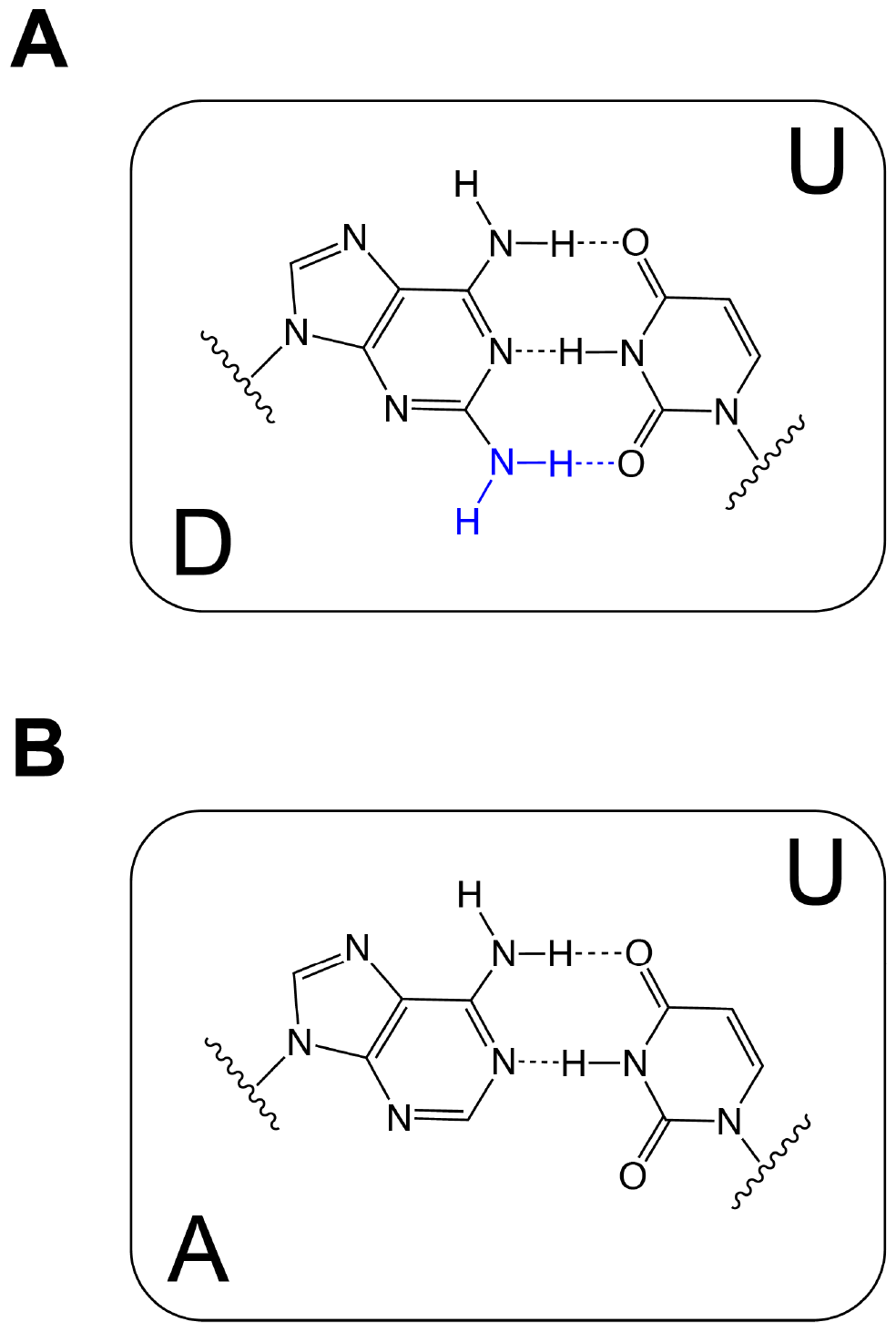
Schematic representation of (A) D:U and (B) A:U base pairs. D:U pair has three hydrogen bonds whereas A:U pair has two.

The potential benefits of D in biological functions have been demonstrated in both RNA and DNA systems, although this substitution has concurrently raised certain concerns. The functionality of RNA with D as a substitution of A is evidenced by the activity of a ribozyme ligase containing only G, D, and U, and the in vitro selection of a variant that contains only D and U.^18^ More recently, it has been shown that replacing ATP with DTP enables an RNA polymerase ribozyme to synthesize longer extension products.^19^ Furthermore, D has been used in antisense oligoribonucleotide therapies owing to its duplex stabilizing ability.^20,21^ In DNA, D can fully substitute for A in certain modern biological systems such as cyanophages.^22–24^ This substitution may confer an evolutionary advantage resulting from the ability to evade genome digestion by host restriction enzymes.^23,24^ Additionally, D-substituted DNA probes demonstrate greater selectivity and stronger hybridization to phage or genomic target DNA.^25^ Consequently, D has been harnessed in various biological applications, including antisense DNA technologies^26^ and gene targeting therapies^27^. However, incorporating D into DNA can undermine the stability of the B-form DNA, sometimes inducing a transition to the Z or A form.^25^ The consequent changes in DNA structure provoke questions regarding the compatibility of D-substituted DNA with the standard DNA-interacting cellular machinery, which might require re-engineering to accommodate D recognition and protein assembly.^28^

Understanding the benefits and challenges of D substitution in RNA and DNA systems has motivated us to evaluate the performance of D in nonenzymatic RNA template copying by reviewing prior studies on this topic. The polymerization of imidazole-activated diaminopurine mononucleotides (ImpD) on a polyU template results in longer oligomers compared to ImpA.^29^ Moreover, nonenzymatic copying on an RNA hairpin containing two templating D residues using activated uridine monomers (ImpU) enhances the elongation rates at lower initial concentrations in the eutectic (water-ice) phase.^30^ Other studies using activated mononucleotides with a different activation group, 2-methyl imidazole (2-MeImp), lead to similar findings. Replacing an A with a D residue in the middle of a 9-mer RNA template increases the efficiency of U incorporation by 3-fold.^10^ Substituting 2-MeImpA with 2-MeImpD as the substrate allows primer extension to proceed at a much lower concentration, particularly in templates with high U content.^31^ Taken together, these studies show that diaminopurine substitution in the template or substrate improves the yield and rate of the nonenzymatic RNA template copying, while leaving the fidelity problem unexplored.

Recent advances in nonenzymatic template copying, particularly the discovery of the prebiotically plausible 2-amino imidazole (2AI) activation group^32^ and the highly reactive 5′-5′ imidazolium-bridged intermediates,^33,34^ warrant a re-evaluation of its efficacy. 2AI-activated mononucleotides readily form bridged dinucleotides^35^ that bind to the template through two Watson-Crick base pairs (Figure S1A) and are the predominant contributors to template-directed primer extension.^33^ Mononucleotides can also react with activated trimers to form monomer-bridged-trimer intermediates with enhanced template affinity and faster rates of primer extension (Figure S1B).^34^ In both scenarios, the 3′-hydroxyl group of the terminal nucleotide of the primer attacks the activated substrate, resulting in the primer being extended by one nucleotide (+1 primer extension) with an activated mononucleotide or an activated trimer as the leaving group (Figure S1). Moreover, the recent development of a deep sequencing protocol, NonEnzymatic RNA Primer Extension Sequencing (NERPE-Seq)^36^, enabled us to not only examine the yield but also the fidelity of the DUCG system in the context of nonenzymatic copying. We have therefore reassessed the effect of diaminopurine in these improved nonenzymatic copying systems.

In this study, we report the enhanced affinity of diaminopurine bridged dinucleotides for a -UU-template. We then compare the rates of nonenzymatic primer extension in the noncanonical DUCG system with the canonical AUCG system. While the D:C mismatch has an unexpectedly elevated reaction rate, the misincorporation of D opposite C hinders the addition of the next base. Moreover, crystallographic studies show that the D:C pair forms only two hydrogen bonds, compared to the three found in the D:U pair. Furthermore, through competition experiments analyzed by next-generation sequencing, we gauge the performance of both the canonical and noncanonical systems under more realistic prebiotic conditions, where four nucleobases co-exist in the non-enzymatic primer extension experiments. Intriguingly, the DUCG system yields a more balanced product base distribution and lowers the mismatch frequency. Moreover, primer extension following mismatches is greatly reduced. Collectively, our findings suggest that diaminopurine improves nonenzymatic template copying, underscoring its potential as a primordial nucleobase.

## Results

### Enhanced binding affinity of D:U pair

Diaminopurine displays superior binding strength to uracil as compared to adenine, presumably due to the additional hydrogen bond and enhanced stacking interactions.^12^ We sought to investigate the effects of this enhanced affinity on the nonenzymatic template copying. To study this, we employed a primer-template-blocker system^34^ with a 2-nt open template region for the bridged-dinucleotide substrate to bind and react (Figure 2A, Table S1). We measured the pseudo-first-order reaction rate constants (k_obs_) for the bridged dinucleotide substrates D^*^D and A^*^A as a function of concentration, and fitted the data using the Michaelis-Menten equation (Figure 2B). The results indicate that D^*^D and A^*^A have similar maximum rates of reaction (k_obs max_), which are 20 h^-1^ and 27 h^-1^, respectively. However, the Michaelis-Menten constant (K_m_) of D^*^D is 20-fold lower than that of A^*^A, 0.033 mM vs 0.64 mM (Figure 2C). This finding underscores the stronger binding of diaminopurine containing substrates and oligonucleotides to RNA templates.

**Figure 2.**
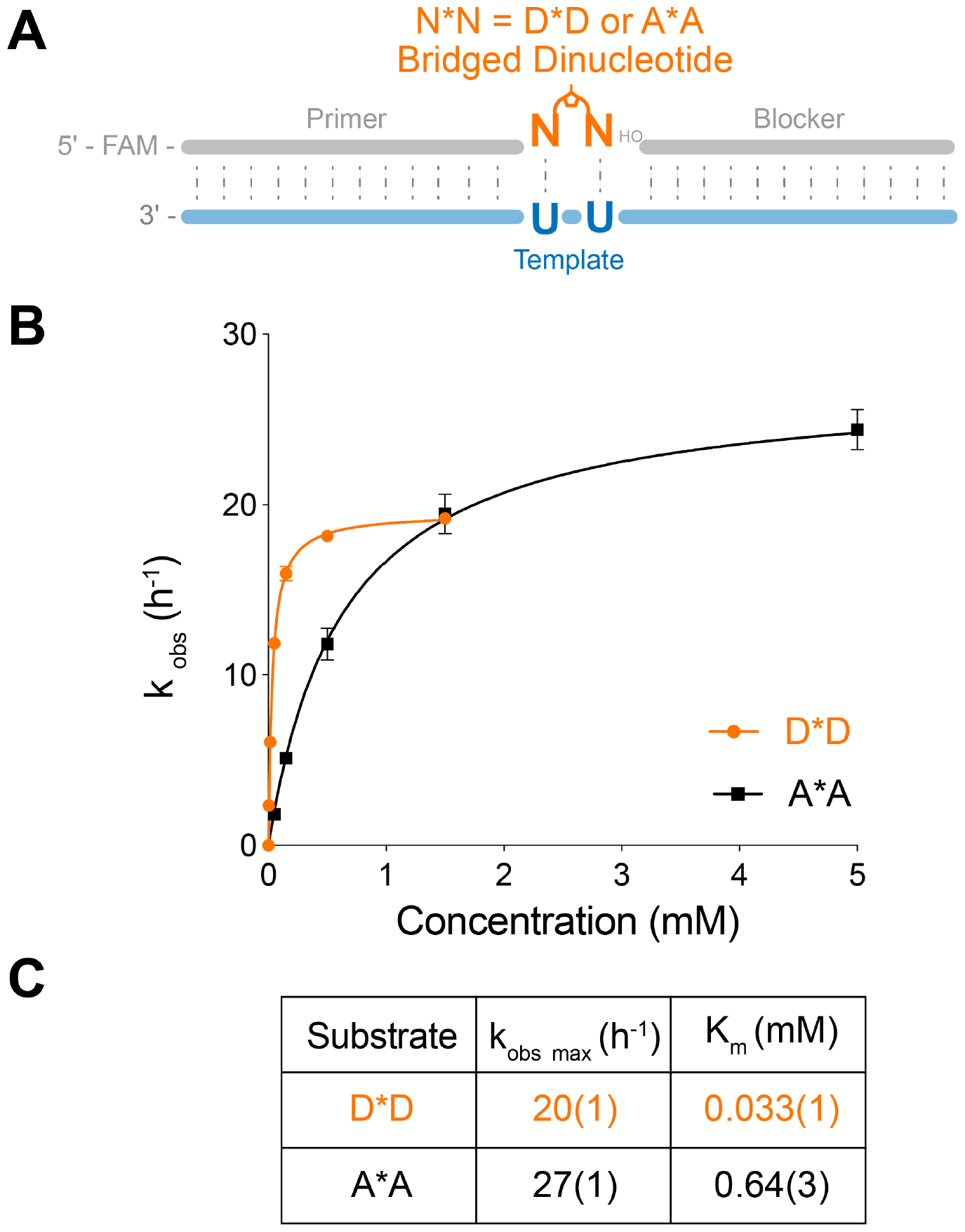
Kinetic study of the bridged diaminopurine and adenine dinucleotides D^*^D and A^*^A. (A) Schematic representation of the nonenzymatic primer extension reactions. (B) Reaction rate vs. concentration curves for D^*^D and A^*^A. (C) Observed maximal rate (k_obs max_) and Michaelis constant (K_m_). Primer extension reactions were performed at the indicated concentrations of bridged dinucleotides (D^*^D or A^*^A), 1.5 µM primer, 2.5 µM template, 3.5 µM blocker, 100 mM MgCl_2_ and 200 mM Tris pH 8.0. Error bars indicate standard deviations of the mean, n=3 replicates. The values of A^*^A is reproduced from refence 34.

The pronounced enhancement of the affinity for D^*^D is in alignment with that predicted by the nearest-neighbor (NN) model. This model predicts the free energy change upon the formation of an RNA double helix, utilizing NN parameters for each stacked pair, a term for the entropy cost of the initial base pairing, and corrections for varied terminal base pairs.^37^ Specifically, the energies associated with D:U stacked pairs and the penalties for the terminal D:U pairs are lower than those of the canonical pairs, resulting in a more negative change in Gibbs free energy value (Table S2).^12,37^ The value of ΔΔG°_37_ predicted by MD/QM modeling, -4.08 ±0.40 kcal/mol, is in qualitative agreement with our experimental observation, where the K_m_ reduction for D^*^D corresponds to a ΔΔG°_25_ of -1.757 kcal/mol (Supplementary Discussion Section). The discrepancy of these values may be attributed to the temperature variations, approximated coaxial stacking and the non-trivial effect of 2-aminoimidazole moieties in bridged dinucleotides, which are modeled as dimers. Despite these differences, both our experimental values and the MD/QM predictions indicate a stronger binding affinity of D:U pairs, likely due to the additional hydrogen bond and enhanced stacking interactions (Table S3).^12^

### Kinetic study of the DUCG system

We examined the impact of substituting D for A on nonenzymatic template-directed primer extension by measuring the rates of primer extension for a complete matrix of 2AI-activated mononucleotides (^*^A, ^*^D, ^*^U, ^*^C, ^*^G) and template nucleotides (A, D, U, C, G) (Figure 3A, Table S4). Because primer extension occurs primarily through an imidazolium-bridged intermediate, we used an activated trinucleotide downstream helper (^*^GAC), which reacts with an activated mononucleotide to form a highly reactive monomer-bridged-trimer intermediate (N^*^GAC) (Figure S1B).^34^ With 20 mM activated monomer and 0.5 mM activated trimer, the template is essentially saturated with N^*^GAC at all times. Although the formation of the bridged intermediate and the subsequent primer extension reactions are two distinct stages within this model system, the first step is relatively fast. Therefore, we could measure pseudo-first-order reaction rate constants to estimate the efficiency of nonenzymatic primer extension. Under these conditions, primer extension with U is slightly faster on a D template than on an A template. Conversely, the incorporation of a D monomer is very slightly faster than that of an A monomer on a U template (Figure 3B).

**Figure 3.**
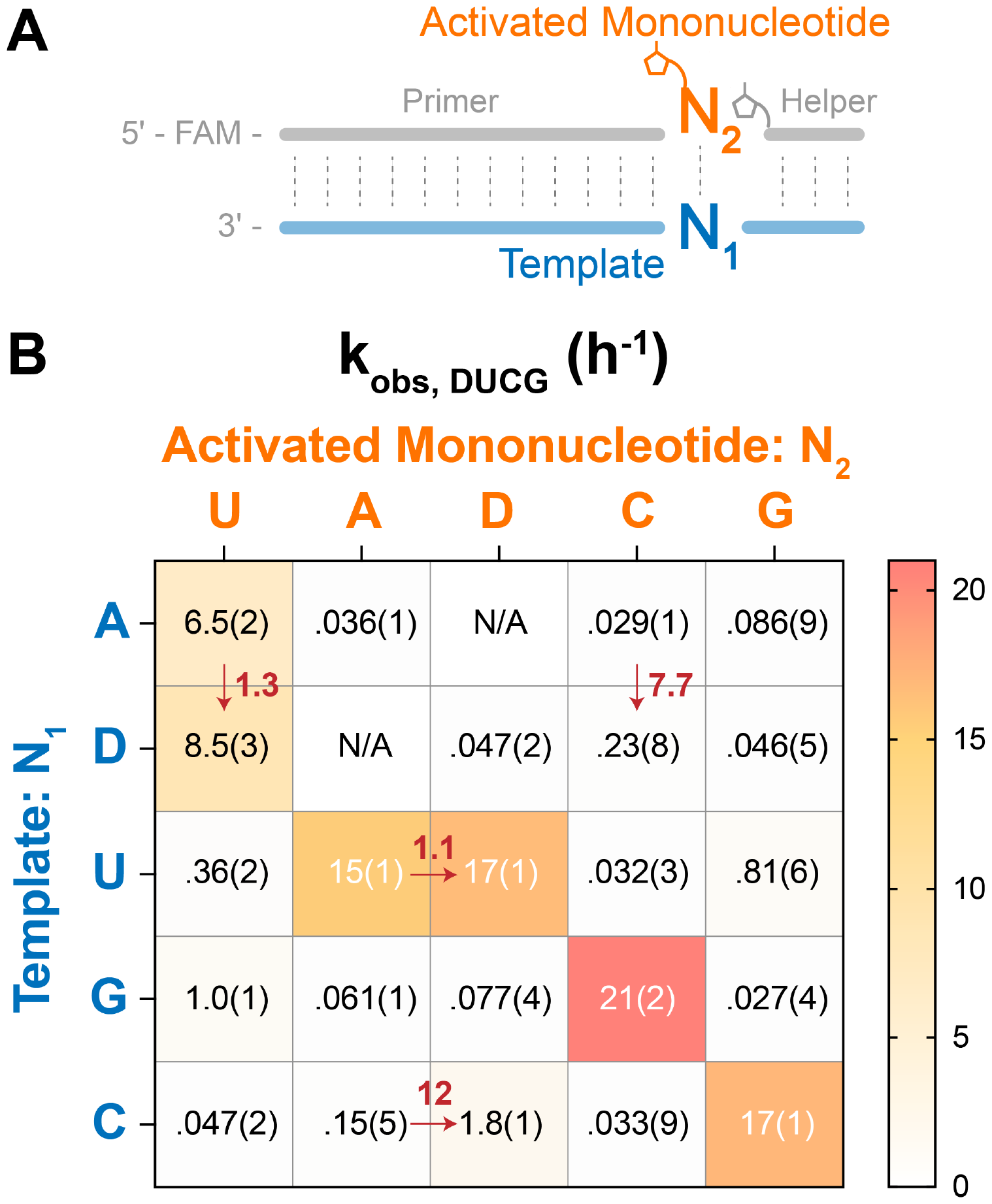
Kinetic study of the DUCG system. (A) Schematic representation of the nonenzymatic primer extension reactions. (B) k_obs_ of the primer extension reactions in different combinations of template: N_1_ and activated mononucleotide: N_2_. Notable rate changes are denoted, with arrows pointing to the direction of these changes. Error bars indicate standard deviations of the mean, n=3 replicates. Primer extension reactions were performed at 20 mM activated mononucleotides, 0.5 mM activated trinucleotide helper, 1.5 µM primer, 2.5 µM template, 100 mM MgCl_2_ and 200 mM Tris pH 8.0. Data of the AUCG systems are adapted from Figure 3 in the reference 38 with permission under a Creative Commons Attribution 4.0 International License. Copyright 2024 American Chemical Society.

Using the above experimental construct, we were also able to measure all mismatched primer extension rates. However, we observed a significant amount of primer-trimer ligation when the N_1_ nucleotide in the template is C. This issue arises because ^*^GAC competes with ^*^N for substrate binding, which interferes with accurate measurement of the rate of mismatched primer extension. To address this problem, we used a modified template with a CUCC overhang and a ^*^AGG trinucleotide helper, to resolve the competitive binding issue with ^*^N (Table S4). We deliberately chose ^*^AGG with a 5′-purine base for a similar stacking interaction between the activated mononucleotide and trinucleotide. This sequence modification enabled more accurate rate measurements of mismatched primer extension. However, it is important to note that the associated downstream template and helper sequences are different from the standard construct in this case.

When we examined the primer extension rates for mismatched base pairs, we were surprised to see that the reaction rates for the D:C mismatch are markedly higher than those of A:C mismatch (12-fold for N_1_:N_2_=C:D and 7.7-fold for N_1_:N_2_=D:C). This increase in reaction rates for D:C pairs raises concerns regarding the fidelity of the DUCG system. To investigate the consequences of enhanced D:C mismatch formation, we measured its stalling effect. In addition, we solved the crystal structures of RNA duplexes containing D:C pairs to help understand their increased formation rate.

### D:C mismatches

#### Stalling effect of D:C mismatches

To address the concern raised by the elevated formation rates of D:C pairs, we investigated the influence of terminal D:C mismatches on subsequent nonenzymatic template primer extension. We prepared template-primer duplexes with either a D:C, D:U or C:G base pair at 3′-end (Table S5) and measured their primer extension rates. We observed that the incorporation of nucleotides following a terminal D:C mismatch is significantly slower than following the complementary base pairs (C:G and D:U) (Figure 4). Therefore, D:C mismatches have a strong stalling effect on downstream primer extension. We quantified this impact using the stalling factor S, defined as the ratio of rates associated with terminal complementary and mismatched base pairs, k_obs_C:G_/k_obs_C:D_ and k_obs_D:U_/k_obs_D:C_. The stalling factors are 5.6 and 12, respectively (Figure 4B, C). This significant stalling effect hinders primer extension after D:C misincorporations and therefore enhances the overall fidelity of template copying in the noncanonical DUCG system.

**Figure 4.**
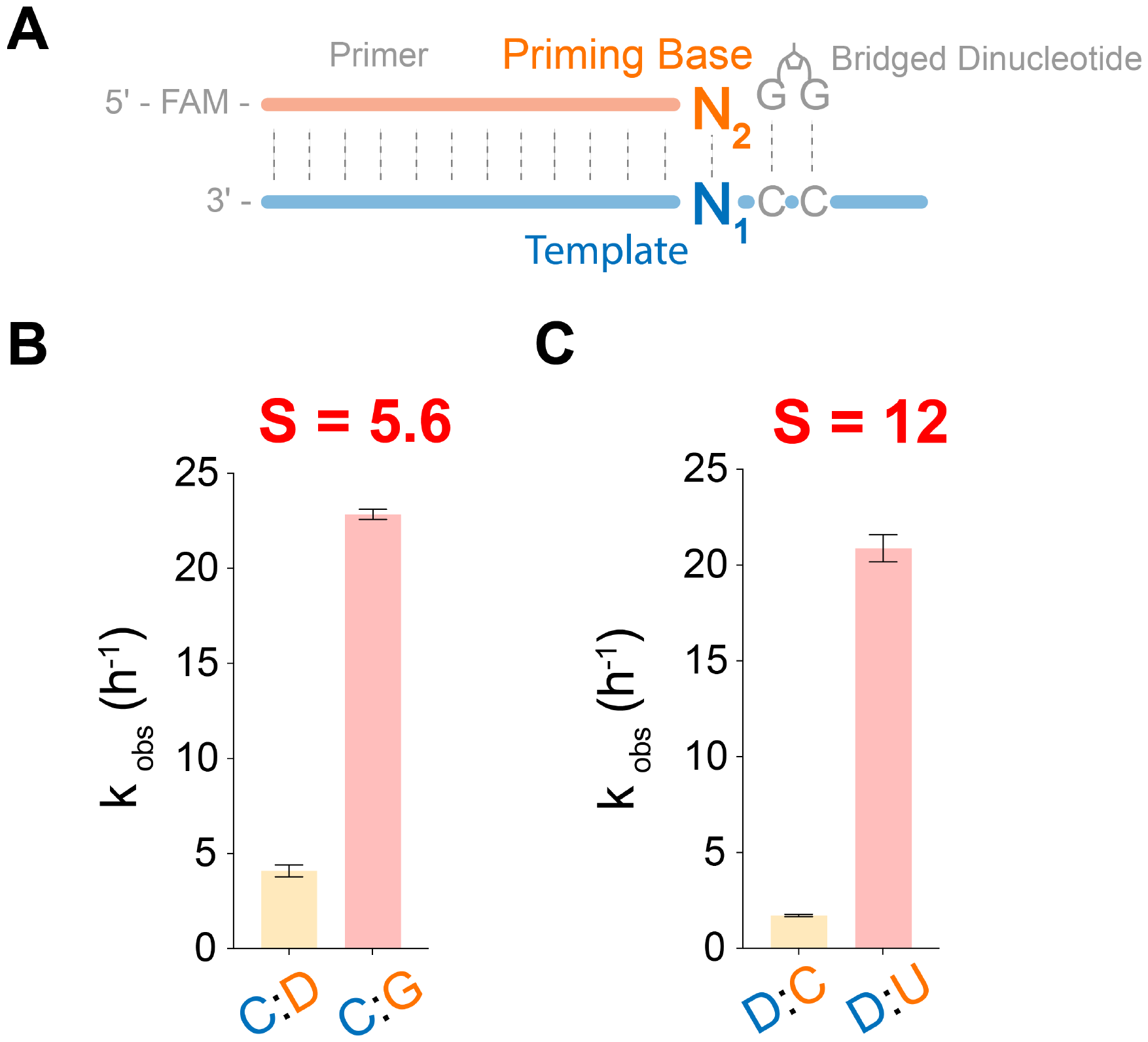
Stalling effects of D:C mismatch. (A) Schematic representation of the primer extension reactions for evaluating the stalling effects of terminal D:C mismatch pairs. (B) Barplot of primer extension reactions (N_1_:N_2_=C:D and C:G). Stalling factor S was calculated from k_obs_C:G_/k_obs_C:D_. (C) Barplot of primer extension reactions (N_1_:N_2_=D:C and D:U). Stalling factor S was calculated from k_obs_D:U_/k_obs_D:C_. Primer extension reactions were performed at 10 mM bridged dinucleotide (G^*^G), 1.5 µM primer, 2.5 µM template, 100 mM MgCl_2_ and 200 mM Tris pH 8.0. Error bars indicate standard deviations of the mean, n=4 replicates.

#### Crystal structures of RNA duplexes containing D:U and D:C pairs

To further understand the structure and properties of the D:U base pair and the D:C mismatch, we designed and synthesized self-complementary RNA sequences that form duplexes with two separated or adjacent D:U or D:C pairs (Table S6). The sequence UD-1, 5′-AGA G<u>D</u>A GAU CU<u>U</u> CUC U-3′, can form two separated D:U pairs with the underlined nucleobases, while the sequence CD-1, 5′-AGA G<u>D</u>A GAU CU<u>C</u> CUC U-3′, can form two separated D:C pairs. Similarly, the sequence UD-2, 5′-AGA GAA G<u>DU</u> CUU CUC U-3′, can form two adjacent D:U pairs with the underlined nucleobases, while the sequence CD-2, 5′-AGA GAA G<u>DC</u> CUU CUC U-3′, can form two adjacent D:C pairs. All four oligonucleotides crystallized within 2-3 days at 20°C under their optimal crystallization conditions (Table S7) and we solved their structures by x-ray diffraction at a resolution higher than 1.65 Å. Data collection and structure refinement statistics are summarized in Table S8 and S9. We found that all four structures adopt the same space group (R32). Each unit cell contains only a single RNA strand, so that each duplex features two identical D-containing base pairs.

Our crystallographic studies show that the D:U base pair has the expected Watson-Crick geometry. Whether separated (UD-1) or adjacent (UD-2) within the RNA duplex, the D:U pairs have identical geometries, and exhibit three hydrogen bonds (Figure 5A). Relative to a canonical A:U base pair, the third hydrogen bond is between the exocyclic 2-amino group and O2 of U. The H-bond distances between N_6_-O_4_, N_1_-N_3_, N_2_-O_2_ in both D:U pairs are identical: 2.8 Å, 2.9 Å and 2.8 Å (Figure 5B, C).

**Figure 5.**
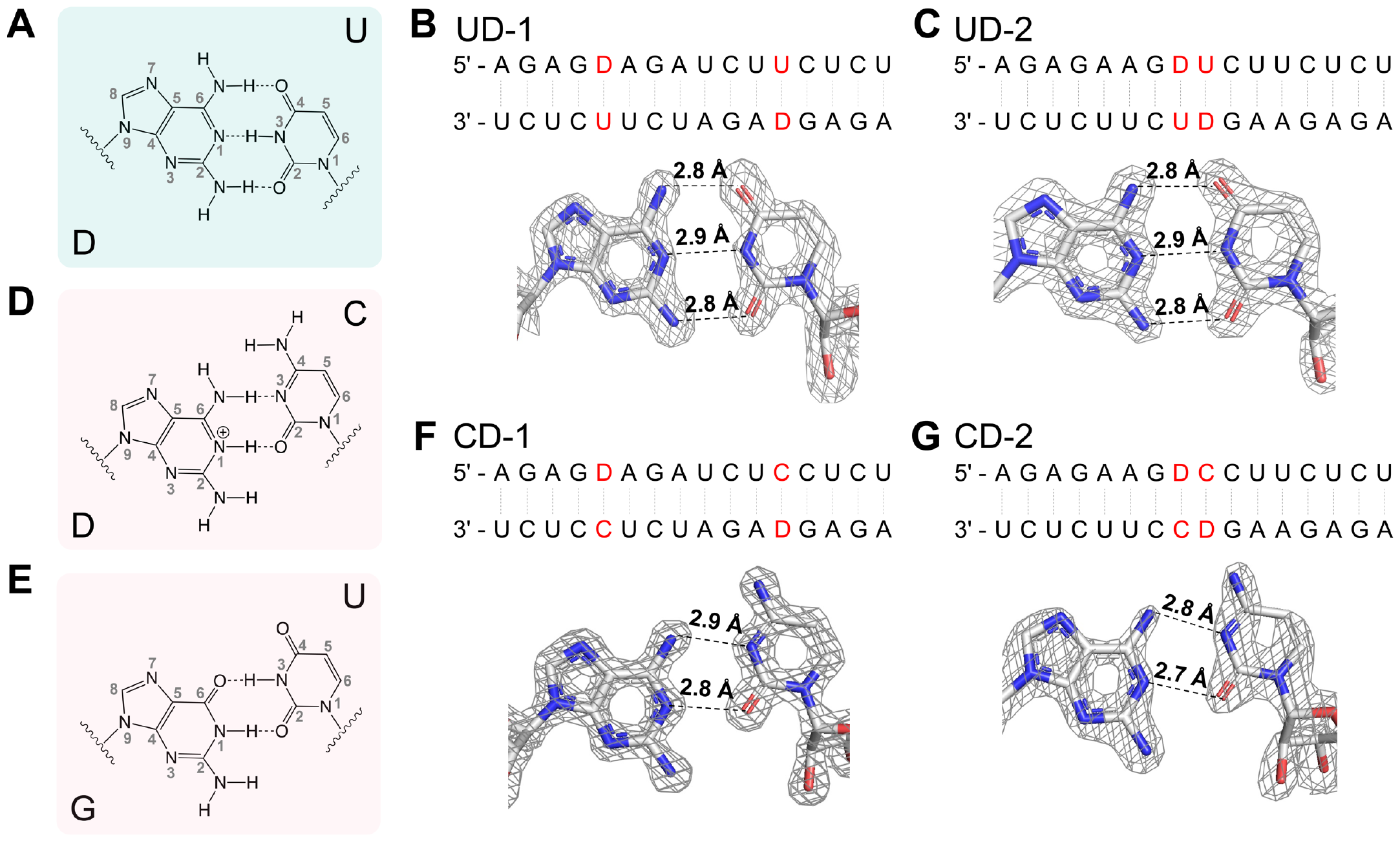
Crystal structures of D:U and D:C pairs. (A) Chemical structure of the complementary D:U pair. (B) Sequences and crystal structures of the UD-1 duplex containing two separated D:U pairs. (C) Sequences and crystal structures of the UD-2 duplex containing two adjacent D:U pairs. (D) Chemical structure of the mismatched D:C pair. (E) Chemical structure of the G:U wobble pair. (F) Sequences and crystal structures of the CD-1 duplex containing two separated D-C pairs. (G) Sequences and crystal structures of the CD-2 duplex containing two adjacent D:C pairs. The grey meshes indicate the corresponding 2Fo-Fc omit maps contoured at 1.5 σ.

Examination of the crystal structures of the CD-1 and CD-2 duplexes shows that the D:C mismatches all adopt the classical wobble base pair geometry that is isomorphic with a G:U wobble pair (Figure 5D, E). The N_6_-N_3_ and N_1_-O_2_ distances are similar for CD-1 and CD-2 duplexes: 2.9 Å vs. 2.8 Å and 2.8 Å vs. 2.7 Å (Figure 5F, G). Based on the observed interatomic distances, the D:C pairs in the CD-1 and CD-2 duplexes appear to have two hydrogen bonds: N_6_-N_3_ and N_1_-O_2._ However, an N_1_-O_2_ hydrogen bond requires N_1_-protonation of D in D:C pairs. Solution-phase NMR studies revealed that the 2-aminopurine (2AP)-cytosine (C) pair exists predominantly as a protonated pair as opposed to an imino tautomer at physiological pH.^39^ Since D has an additional electron-donating amino group, N_1_ in D has a higher pK_a_ value and is more likely to be protonated than N_1_ in 2AP.^40^ Therefore, D in the D:C pairs likely exists as a protonated D in the canonical amino tautomeric form (Figure 5D).

### Sequencing analysis of the AUCG (canonical) and DUCG (noncanonical) systems

We next conducted competition experiments, in which all four activated monomers were used to copy a random sequence template region. We then used deep RNA sequencing to compare the efficiency and fidelity of the non-canonical DUCG system to the canonical AUCG system in nonenzymatic primer extension. We adapted the protocol for NonEnzymatic RNA Primer Extension Sequencing (NERPE-Seq)^36^: two sets of mixed bridged dinucleotides (N^*^N, where N=A, U, C, G in the AUCG system and N=D, U, C, G in the DUCG system) were added to the respective self-priming hairpin constructs (Table S10) with a 6-nucleotide long randomized region containing all four bases (6N, where N=A, U, C, G in the AUCG system and N=D, U, C, G in the DUCG system). The reaction mixtures were incubated for 24 hours then quenched and processed for next-generation sequencing (Figure 6A). We used Moloney Murine Leukemia Virus (MMLV) Reverse Transcriptase for cDNA synthesis, which has been reported to incorporate T opposite D with high fidelity.^41^ We then examined the resulting sequences to determine the yield, fidelity, product distribution and mismatch patterns of the products of nonenzymatic copying.

**Figure 6.**
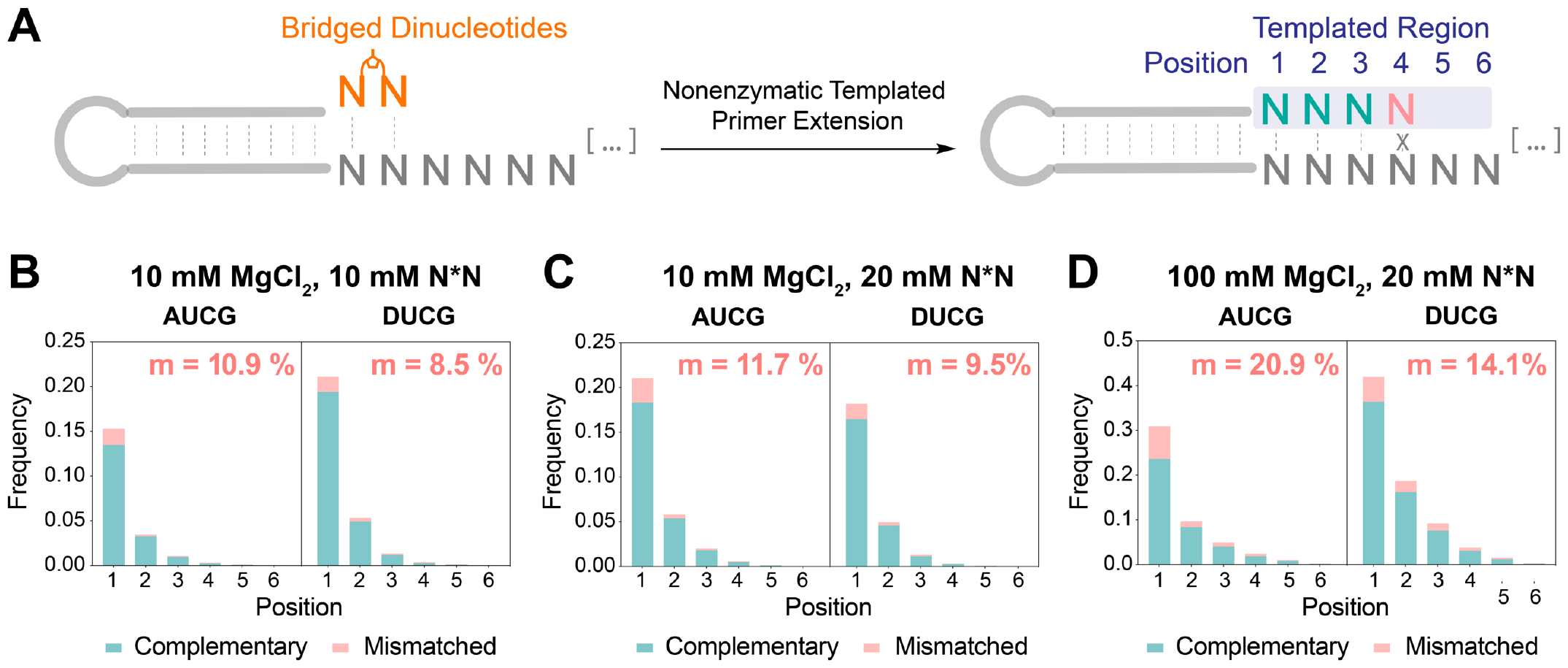
(A) Schematic representation of primer extension on a hairpin primer/template. (B)-(D) Yield and fidelity stacked barplots show the position-dependent complementary and mismatched incorporation frequency in AUCG and DUCG systems. At each position, the sum of the frequencies of primer extension with a complementary nucleotide, a mismatched nucleotide and no extension was normalized to 1. Mismatch frequency (m) is calculated for the extended sequence at position 1-4. Stacked barplots are generated for the following reaction conditions: (B) 10 mM MgCl_2_ and 10 mM N^*^N (C) 10 mM MgCl_2_ and 20 mM N^*^N in (D) 100 mM MgCl_2_ and 20 mM N^*^N. Reactions conditions: 1 μM hairpin, 1.2 μM 5′ Handle Block, 200 mM Tris pH 8.0, 100 mM/10 mM MgCl_2_, 20 mM/10 mM N^*^N, incubated at RT for 24h.

#### Yield and fidelity

We determined the extent and fidelity of template-directed primer extension following a single addition of activated substrates to the primer/template hairpin construct. We used three different reaction conditions to investigate the influence of MgCl_2_ and bridged dinucleotide (N^*^N) concentration: 10 mM MgCl_2_, 10 mM N^*^N; 10 mM MgCl_2_, 20 mM N^*^N; 100 mM MgCl_2_, 20 mM N^*^N. Each individual product sequence was categorized as complementary, mismatched, or unextended. We then generated stacked barplots with position-dependent frequencies of complementary and mismatched incorporation. Mismatch frequency (m) was computed as the fraction of mismatched over total incorporations across all positions.

Our comparative analysis reveals that the DUCG system exhibits a modest improvement over the canonical AUCG system in both yield and fidelity. The DUCG system exhibits a higher frequency of position-dependent incorporation under most tested reaction conditions (Figure 6B-D). The system also has a higher frequency of total extended products: a 38% increase in 10 mM MgCl_2_, 10 mM N^*^N, and a 36% increase in 100 mM MgCl_2_, 20 mM N^*^N (Table S11). Furthermore, an improvement in fidelity is observed across all reactions conditions in the DUCG system, with a notable 20-30% decrease in error rate compared to the AUCG system (Figure 6B-D). The fidelity advantage of the DUCG system is most apparent at high MgCl_2_ concentration, which catalyzes the hydrolysis of bridged dinucleotides, thereby reducing the bridged-to-mononucleotide ratio and consequently decreasing fidelity.^5^ In addition, the mismatch ratio is not affected by the concentration of bridged dinucleotides, as the bridged-to-mononucleotide ratio is independent of substrate concentration. Overall, the elevated yield and increased fidelity observed in the DUCG system affirm its advantage in nonenzymatic template copying.

#### Distribution of product bases

After probing the yield and fidelity of the AUCG and DUCG systems, we next examined the product composition. Nonenzymatic primer extension products generated by the addition of an equimolar mixture of all four activated canonical nucleotides tend to be rich in G and C, in part because the A:U base pair is weaker than the G:C pair.^5^ We wondered whether the DUCG system could alleviate this biased incorporation. We quantified the product base distribution among fully complementary products for both the AUCG and the DUCG systems (Figure 7). To better compare the two systems, we also plotted the ratio of frequencies between the DUCG and the AUCG systems. Numbers above 1 (colored in red) indicate enrichment and those below 1 (colored in blue) indicate diminishment in the DUCG system. The product base distribution of the AUCG system across all reaction conditions is heavily biased towards G and C incorporation, aligning with findings from prior research.^5^ In contrast, the DUCG system has a more even product base distribution with an significant increase in the incorporation of both D and U. For example, under the 10 mM MgCl_2_ and 10 mM N^*^N condition, product base frequency of D, compared to A, increased from 0.10 to 0.17 and that of U increased from 0.06 to 0.10 in position 1. This phenomenon is especially significant at lower MgCl_2_ concentration as the effect of enhanced affinity of the D:U base pair becomes more dominant. Overall, the DUCG system effectively mitigates the issue of biased incorporation of G and C across all reaction conditions.

**Figure 7.**
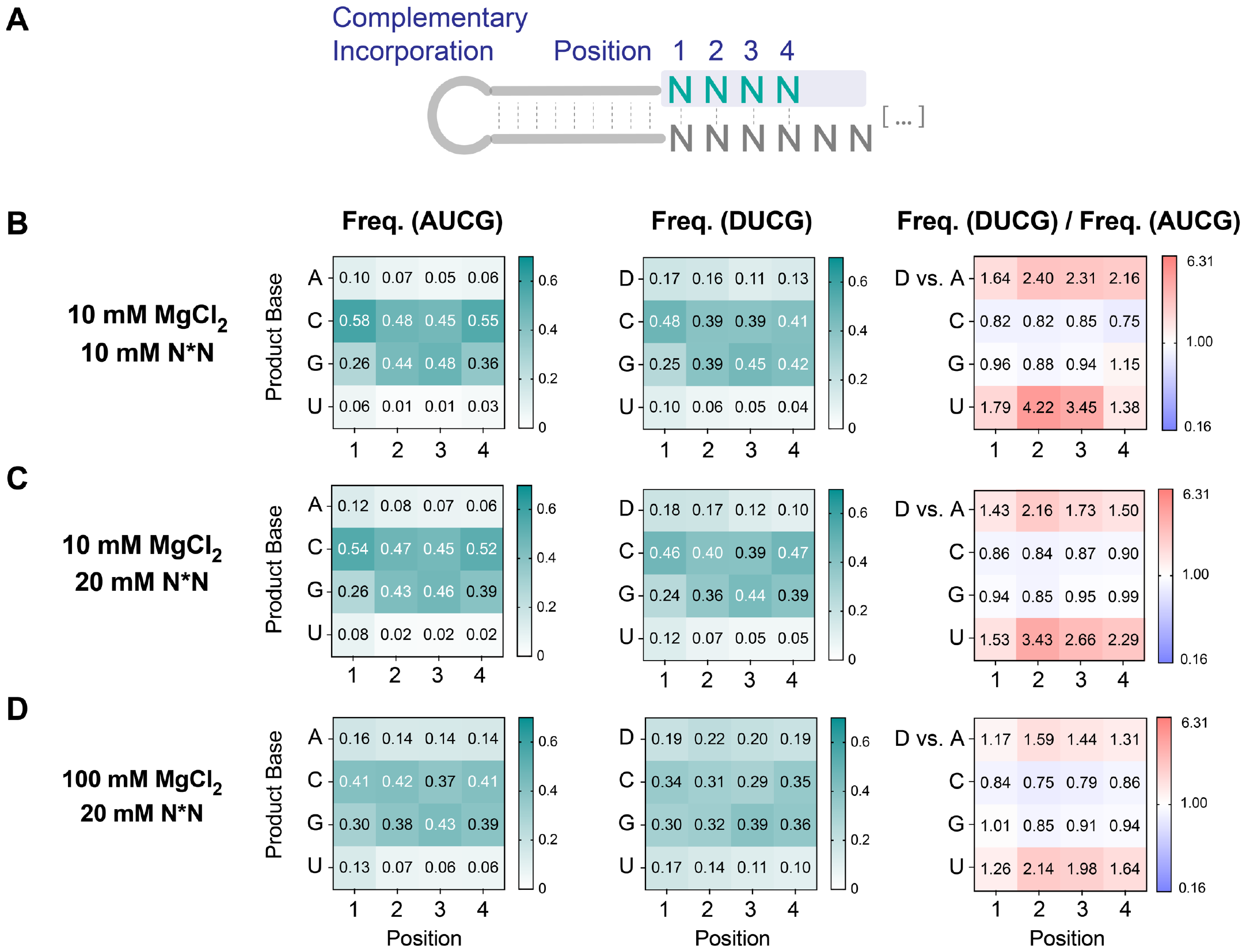
DUCG system decreases biases in complementary product incorporation by enriching D and U in the product base distribution. (A) Schematic representation of the product base distribution. (B)-(D) Position-dependent product base frequency in the AUCG & DUCG systems and the frequency ratio between AUCG and DUCG. Heatmaps are generated for the following reaction conditions: (B) 10 mM MgCl_2_ and 10 mM N^*^N (C) 10 mM MgCl_2_ and 20 mM N^*^N (D) 100 mM MgCl_2_ and 20 mM N^*^N. For the frequency ratio heatmap, red represents greater frequency in the DUCG system whereas blue represents greater frequency in the AUCG system.

Beyond examining the product base distribution, we delved into the distribution of inferred bridged dinucleotides, which serve as the primary substrates for template copying. Leveraging the template composition information made possible by deep sequencing, we deduced the normalized distribution of the 16 possible bridged dinucleotides involved in the nonenzymatic primer extension. In alignment with previous research,^5^ the AUCG system exhibited an increased frequency of G and C in both the first and second positions of the inferred bridged dinucleotides (Figure S2), attributed to their enhanced binding affinity with the template. We were gratified to see that the DUCG system substantially mitigated this bias by enhancing the incorporation of D^*^N and U^*^N across all reaction conditions (Figure S2B-D). The increased incorporation frequency observed with D^*^N and U^*^N is most pronounced when the second nucleotide is also a D or U (i.e., D^*^D, D^*^U, U^*^D, and U^*^U). This outcome underscores the impact of the enhanced binding affinity of diaminopurine on the bridged dinucleotide primer extension pathway (Figure S1A). It serves as an effective mechanism for equalizing the product distribution in nonenzymatic primer extension processes, thereby maximizing the diversity of inherited genetic information.

#### Mismatch composition and stalling

We measured the position-dependent frequency of all 12 possible mismatches in the AUCG and DUCG systems. Consistent with previous research,^5^ the A:G and D:G (template: product) pair is the most frequent mismatch at position 1 in both systems across all reaction conditions (Figure S3). Interestingly, the D:C and C:D mismatches are not significantly overrepresented, when compared with A:C and C:A mismatches.

The deep sequencing data also allowed us to look at the effect of mismatches on subsequent primer extension. In both systems, the probability of primer extension past a mismatched pair at position 1 is significantly lower compared to that past a complementary pair (Figure 8A). The complementary pairs exhibit a 4 to 9-fold higher likelihood of extension compared to mismatched pairs under all tested reaction conditions. This indicates that once a mismatch is incorporated, it is less likely to extend further than a matched nucleotide, as expected from the stalling effect we observed previously for D:C mismatch pairs. Furthermore, we quantified the extension probability over each type of mismatch at the +1 position (Figure S4A) along with the corresponding stalling factor (Figure S4B). The stalling factors range from roughly 2 to 10 depending on the mismatch.

**Figure 8.**
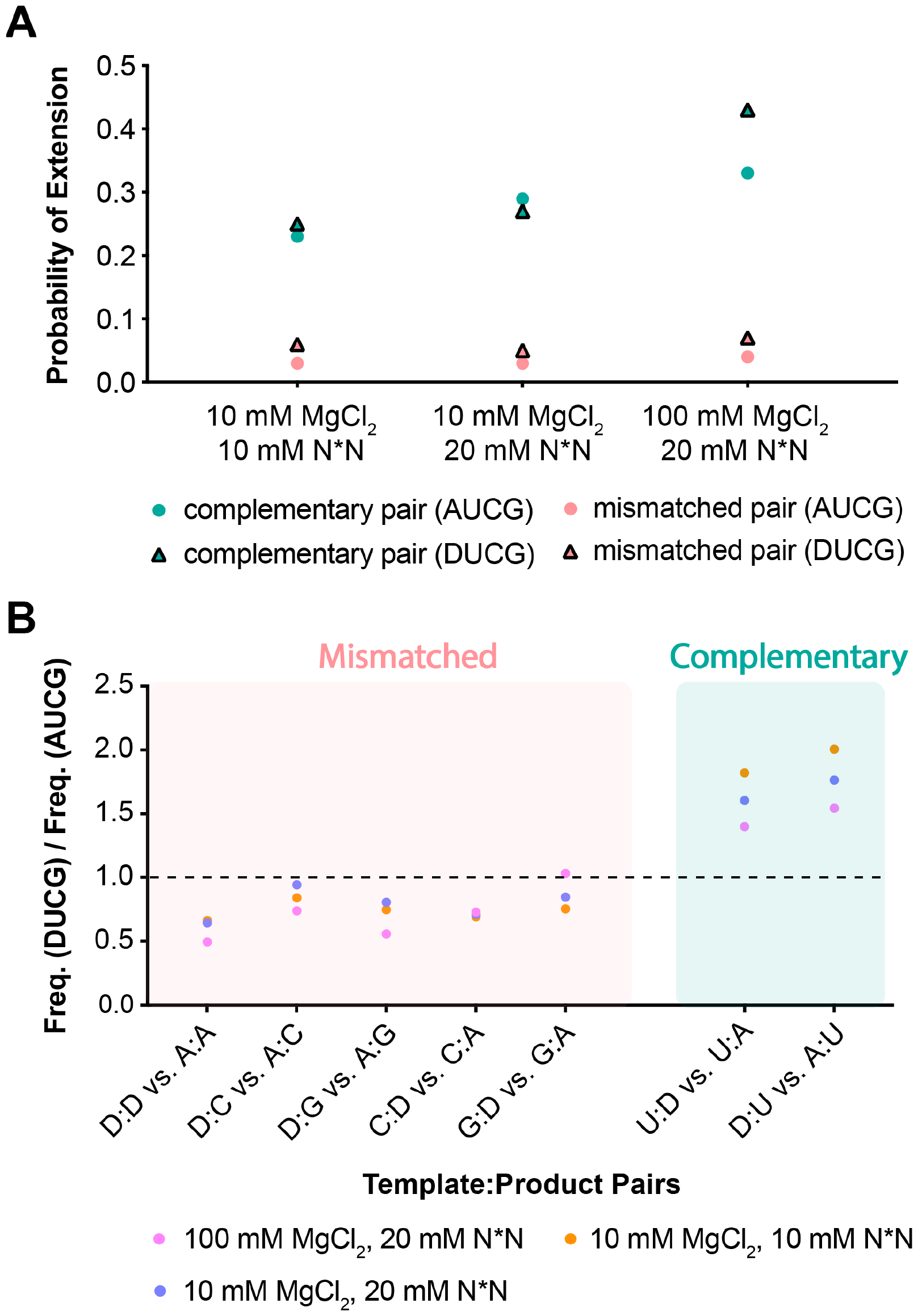
The effect of mismatches on subsequent primer extension and the impact of diaminopurine substitution. (A) Probability of extension following a complementary pair versus a mismatched pair at position 1 in the AUCG and DUCG systems. (B) Frequency ratio of overall T:P pairs distribution among all incorporation (mismatched incorporation + complementary incorporation) between the DUCG and AUCG systems. Frequency ratios of T:P pairs containing D are selected and plotted.

The error frequency for primer extension after a mismatch is also significantly higher than the overall mismatch frequency, in both systems and across all reaction conditions (Figure S5). The mismatched priming base pair seems to lack the proper conformation required to enforce correct base pairing on the subsequent nucleotide incorporation. Overall, all template:product (T:P) pairs containing D display a decrease in subsequent mismatched pair frequency and a corresponding increase in complementary frequency (Figure 8B), underscoring the advantage of the DUCG system in the fidelity of nonenzymatic template copying.

## Discussion

To ensure the efficient transmission of genetic information before the advent of ribozymes, we examined modifications of the A:U base pair that would address the issue of incorporation biases in nonenzymatic RNA template copying. Our previous study demonstrated that the A:s^2^U (2-thio-U) pair stabilizes RNA duplexes by reducing the desolvation penalty and pre-organizing single-stranded RNA during hybridization.^38^ In this study, we evaluated the alternative D:U base pair, which enhances binding through an additional hydrogen bond and stronger stacking interactions. As a result, we observed a 20-fold increase in the binding of the substrate (D^*^D versus A^*^A) to a -UU-template (Figure 2). Accordingly, the advantage of diaminopurine is less pronounced at higher substrate concentrations (Figure 2 and 3) and in the presence of a downstream activated trimer helper (Figure 3). The latter reacts with an activated mononucleotide to form a highly reactive monomer-bridged-trimer intermediate^34^, masking the advantage of diaminopurine’s increased binding strength. These results collectively suggest that the distinctive benefit of diaminopurine is most notable at low and sub-saturating substrate concentrations.

We examined the diaminopurine system in both a defined construct, and in competition experiments where all four nucleotides vied for incorporation on a mixed-sequence template. The most notable difference is the drastic rate increase for C:D (12-fold) and D:C (7.7-fold) mismatches in the defined construct (Figure 3B), in contrast to their marginal (Figure S3) or reduced (Figure 8B) presence in the competition experiments. Crystallographic studies reveal that the protonated amino form of D can form a wobble-type base pair with C via two hydrogen bonds (Figure 5), which may account for the increased rate of D:C mismatches in the defined construct (Figure 3B). In competition experiments, D:C mismatches are likely to be outcompeted by the correct C:G and D:U pairs during incorporation, accounting for their infrequent occurrence (Figure S3). Additionally, the strong stalling effects of the D:C mismatches (Figure 4, Figure S4B) further alleviate concerns regarding the fidelity of the DUCG system.

In addition to the inhibitory effect of the D:C mismatches on subsequent primer extension, our competition experiments showed a pronounced stalling effect across all other mismatched pairs, effectively hindering downstream primer extension and therefore improving the system’s overall fidelity.^42^ The A:G and D:G (template: product) mismatches emerged as the most significant among all 12 mismatches studied (Figure S3). The higher prevalence of A:G and D:G mismatches as compared to G:A and G:D mismatches most likely stems from the effective G template depletion by the correct C incorporation.^5^ We were pleased to find that the A:G and D:G mismatches have the lowest probability of extension (Figure S4A) and the highest stalling factor (Figure S4B). Furthermore, mismatches that extend are more susceptible to further misincorporations (Figure S5), likely due to the lack of conformational rigidity in the priming base pair. Overall, the probability of extension over a mismatched pair is much lower than that of a complementary pair in both systems (Figure 8A), effectively reducing downstream error propagation and enhancing overall fidelity.

Through deep sequencing, we observed that the DUCG system outperforms the AUCG system in both yield and fidelity (Figure 6). This effect is most notable at low MgCl_2_ and substrate concentrations, which are more compatible with the fatty acid based membranes^43^ and which may be more prebiotically plausible^44,45^. Moreover, the copying of template nucleotides is more uniform in the DUCG system under all tested reaction conditions (Figure 7, Figure S2). Despite improvements, the G and C-bias issue still persists (Figure 7), since the D:U base-pair is not as strong as the G:C base-pair. This is likely attributable to the repulsive electrostatic cross-interactions between the functional groups in the D:U pairs^11^, as supported by thermodynamic^13,46^ and computational studies^12^.

## Conclusions

In summary, diaminopurine (D) has emerged as a promising primordial nucleobase candidate, showing advantageous attributes in nonenzymatic primer extension within the DUCG system. With an extra hydrogen bond and increased stacking interactions, D:U pairs exhibit enhanced copying at low substrate concentrations, as reflected by a significant reduction in K_m_. Furthermore, while D:C mismatches are significant in a defined construct, they do not compromise the system’s fidelity in a competition setting. Moreover, they, along with other mismatches, demonstrate a strong stalling effect on primer extension. Thus, the DUCG system outperforms the canonical AUCG system in both yield and fidelity, especially at lower MgCl_2_ and substrate concentrations. We may therefore ask whether the DUCG system is a plausible primordial genetic alphabet. The answer to this question depends primarily on whether a plausible high yielding prebiotic synthesis of diaminopurine nucleotides can be discovered. A second key question is whether the greater RNA duplex stability that results from replacing A with D is an advantage or a disadvantage in terms of nonenzymatic RNA replication. Further research into the DUCG and other potential primordial genetic alphabets will be required to address these questions.

## ASSOCIATED CONTENT

### Supporting Information

The Supporting Information is available free of charge on the ACS Publications website.

The raw sequencing files, processing worksheets and code can be found in the OSF.io repository: https://osf.io/zscy8/.

Material and Methods; Supplementary Discussion; Supplementary Figures and Tables (PDF)

## Author Contributions

The manuscript was written through the contributions of all authors.

## Notes

The authors declare no competing financial interest.

## ACKNOWLEDGMENT

J.W.S. is an Investigator of the Howard Hughes Medical Institute. This work was supported in part by grants from the NSF (2104708), the Sloan Foundation (G-2022-19518) and the Gordon and Betty Moore Foundation (11479) to J.W.S. This work was also supported by the University of Pennsylvania to L.Z.

The authors thank Dr. Daniel Duzdevich for preliminary sequencing experiments, advice on the sequencing assay and result interpretation, and comments on the manuscript. The authors also thank Dr. Marco Todisco for helpful discussion on NN predictions. In addition, we thank Ulandt Kim and other staff of the MGH Department of Molecular biology Next-Generation Sequencing Core for sample validation and MiSeq runs. In addition, we thank the staff at the Advanced Light Source (ALS) beamline 201.

The Berkeley Center for Structural Biology is supported in part by the Howard Hughes Medical Institute. The Advanced Light Source is a Department of Energy Office of Science User Facility under Contract No. DE-AC02-05CH11231. The ALS-ENABLE beamlines are supported in part by the National Institutes of Health, National Institute of General Medical Sciences, grant P30 GM124169.

## ABBREVIATIONS

D: diaminopurine
NN: nearest-neighbor
MD/QM: molecular dynamics/quantum mechanics

## REFERENCES

(1) Szostak, J. W.; Bartel, D. P.; Luisi, P. L. Synthesizing Life. Nature 2001, 409 (6818), 387–390.

(2) Orgel, L. E. Evolution of the Genetic Apparatus. Journal of Molecular Biology 1968, 38 (3), 381–393.

(3) Gilbert, W. Origin of Life: The RNA World. Nature 1986, 319 (6055), 618–618.

(4) Szostak, J. W. The Eightfold Path to Non-Enzymatic RNA Replication. Journal of Systems Chemistry 2012, 3 (1), 2.

(5) Duzdevich, D.; Carr, C. E.; Ding, D.; Zhang, S. J.; Walton, T. S.; Szostak, J. W. Competition between Bridged Dinucleotides and Activated Mononucleotides Determines the Error Frequency of Nonenzymatic RNA Primer Extension. Nucleic Acids Research 2021, 49 (7), 3681–3691.

(6) Callahan, M. P.; Smith, K. E.; Cleaves, H. J.; Ruzicka, J.; Stern, J. C.; Glavin, D. P.; House, C. H.; Dworkin, J. P. Carbonaceous Meteorites Contain a Wide Range of Extraterrestrial Nucleobases. Proceedings of the National Academy of Sciences 2011, 108 (34), 13995–13998.

(7) Szabla, R.; Zdrowowicz, M.; Spisz, P.; Green, N. J.; Stadlbauer, P.; Kruse, H.; Šponer, J.; Rak, J. 2,6-Diaminopurine Promotes Repair of DNA Lesions under Prebiotic Conditions. Nat Commun 2021, 12 (1), 3018.

(8) Kim, H.-J.; Benner, S. A. Prebiotic Stereoselective Synthesis of Purine and Noncanonical Pyrimidine Nucleotide from Nucleobases and Phosphorylated Carbohydrates. Proceedings of the National Academy of Sciences 2017, 114 (43), 11315–11320.

(9) Caldero-Rodríguez, N. E.; Ortiz-Rodríguez, L. A.; Gonzalez, A. A.; Crespo-Hernández, C. E. Photostability of 2,6-Diaminopurine and Its 2′-Deoxyriboside Investigated by Femtosecond Transient Absorption Spectroscopy. Phys. Chem. Chem. Phys. 2022, 24 (7), 4204–4211.

(10) Kozlov, I. A.; Orgel, L. E. Nonenzymatic Oligomerization Reactions on Templates Containing Inosinic Acid or Diaminopurine Nucleotide Residues. Helvetica Chimica Acta 1999, 82 (11), 1799–1805.

(11) Hartel, C.; Göbel, M. W. Substitution of Adenine by Purine-2,6-Diamine Improves the Nonenzymatic Oligomerization of Ribonucleotides on Templates Containing Thymidine. Helvetica Chimica Acta 2000, 83 (9), 2541–2549.

(12) Hopfinger, M. C.; Kirkpatrick, C. C.; Znosko, B. M. Predictions and Analyses of RNA Nearest Neighbor Parameters for Modified Nucleotides. Nucleic Acids Research 2020, 48 (16), 8901–8913.

(13) Howard, F. B.; Frazier, J.; Miles, H. T. A New Polynucleotide Complex Stabilized by 3 Interbase Hydrogen Bonds, Poly-2-Aminoadenylic Acid + Polyuridylic Acid. J Biol Chem 1966, 241 (18), 4293–4295.

(14) Bailly, C.; Waring, M. J. The Use of Diaminopurine to Investigate Structural Properties of Nucleic Acids and Molecular Recognition between Ligands and DNA. Nucleic Acids Research 1998, 26 (19), 4309–4314.

(15) Cristofalo, M.; Kovari, D.; Corti, R.; Salerno, D.; Cassina, V.; Dunlap, D.; Mantegazza, F. Nanomechanics of Diaminopurine-Substituted DNA. Biophys J 2019, 116 (5), 760–771.

(16) Wu, X.; Delgado, G.; Krishnamurthy, R.; Eschenmoser, A. 2,6-Diaminopurine in TNA: Effect on Duplex Stabilities and on the Efficiency of Template-Controlled Ligations1. Org. Lett. 2002, 4 (8), 1283–1286.

(17) Zhou, L.; Ding, D.; Szostak, J. W. The Virtual Circular Genome Model for Primordial RNA Replication. RNA 2021, 27 (1), 1–11.

(18) Reader, J. S.; Joyce, G. F. A Ribozyme Composed of Only Two Different Nucleotides. Nature 2002, 420 (6917), 841–844.

(19) Attwater, J.; Tagami, S.; Kimoto, M.; Butler, K.; T. Kool, E.; Wengel, J.; Herdewijn, P.; Hirao, I.; Holliger, P. Chemical Fidelity of an RNA Polymerase Ribozyme. Chemical Science 2013, 4 (7), 2804–2814.

(20) Lamm, G. M.; Blencowe, B. J.; Sparoat, B. S.; Iribarren, A. M.; Ryder, U.; Lamond, A. I. Antisense Probes Containing 2-Aminoadenosine Allow Efficient Depletion of U5 snRNP from HeLa Splicing Extracts. Nucleic Acids Research 1991, 19 (12), 3193–3198.

(21) Barabino, S. M. L.; Sproat, B. S.; Lamond, A. I. Antisense Probes Targeted to an Internal Domain in U2 snRNP Specifically Inhibit the Second Step of Pre-mRNA Splicing. Nucleic Acids Research 1992, 20 (17), 4457–4464.

(22) Kirnos, M. D.; Khudyakov, I. Y.; Alexandrushkina, N. I.; Vanyushin, B. F. 2-Aminoadenine Is an Adenine Substituting for a Base in S-2L Cyanophage DNA. Nature 1977, 270 (5635), 369–370.

(23) Zhou, Y.; Xu, X.; Wei, Y.; Cheng, Y.; Guo, Y.; Khudyakov, I.; Liu, F.; He, P.; Song, Z.; Li, Z.; Gao, Y.; Ang, E. L.; Zhao, H.; Zhang, Y.; Zhao, S. A Widespread Pathway for Substitution of Adenine by Diaminopurine in Phage Genomes. Science 2021, 372 (6541), 512–516.

(24) Pezo, V.; Jaziri, F.; Bourguignon, P.-Y.; Louis, D.; Jacobs-Sera, D.; Rozenski, J.; Pochet, S.; Herdewijn, P.; Hatfull, G. F.; Kaminski, P.-A.; Marliere, P. Noncanonical DNA Polymerization by Aminoadenine-Based Siphoviruses. Science 2021, 372 (6541), 520–524.

(25) Chollet, A.; Kawashima, E. DNA Containing the Base Analogue 2-Aminoadenine: Preparation, Use as Hybridization Probes and Cleavage by Restriction Endonucleases. Nucleic Acids Research 1988, 16 (1), 305–317.

(26) Rosenbohm, C.; Pedersen, D. S.; Frieden, M.; Jensen, F. R.; Arent, S.; Larsen, S.; Koch, T. LNA Guanine and 2,6-Diaminopurine. Synthesis, Characterization and Hybridization Properties of LNA 2,6-Diaminopurine Containing Oligonucleotides. Bioorganic & Medicinal Chemistry 2004, 12 (9), 2385–2396.

(27) Gao, S.; Guan, H.; Bloomer, H.; Wich, D.; Song, D.; Khirallah, J.; Ye, Z.; Zhao, Y.; Chen, M.; Xu, C.; Liu, L.; Xu, Q. Harnessing Non-Watson–Crick’s Base Pairing to Enhance CRISPR Effectors Cleavage Activities and Enable Gene Editing in Mammalian Cells. Proceedings of the National Academy of Sciences 2024, 121 (2), e2308415120.

(28) Grome, M. W.; Isaacs, F. J. ZTCG: Viruses Expand the Genetic Alphabet. Science 2021, 372 (6541), 460–461.

(29) Webb, T. R.; Orgel, L. E. Template Directed Reactions of 2-Aminoadenylic Acid Derivatives. Nucleic Acids Research 1982, 10 (14), 4413–4422.

(30) Monnard, P.-A.; Szostak, J. W. Metal-Ion Catalyzed Polymerization in the Eutectic Phase in Water-Ice: A Possible Approach to Template-Directed RNA Polymerization. J Inorg Biochem 2008, 102 (5–6), 1104–1111.

(31) Grzeskowiak, K.; Webb, T. R.; Orgel, L. E. Template-Directed Synthesis with 2-Aminoadenosine. J Mol Evol 1984, 21 (1), 81–83.

(32) Li, L.; Prywes, N.; Tam, C. P.; O’Flaherty, D. K.; Lelyveld, V. S.; Izgu, E. C.; Pal, A.; Szostak, J. W. Enhanced Nonenzymatic RNA Copying with 2-Aminoimidazole Activated Nucleotides. J. Am. Chem. Soc. 2017, 139 (5), 1810–1813.

(33) Walton, T.; Zhang, W.; Li, L.; Tam, C. P.; Szostak, J. W. The Mechanism of Nonenzymatic Template Copying with Imidazole-Activated Nucleotides. Angewandte Chemie International Edition 2019, 58 (32), 10812–10819.

(34) Ding, D.; Zhou, L.; Giurgiu, C.; Szostak, J. W. Kinetic Explanations for the Sequence Biases Observed in the Nonenzymatic Copying of RNA Templates. Nucleic Acids Res 2022, 50 (1), 35–45.

(35) Walton, T.; Szostak, J. W. A Kinetic Model of Nonenzymatic RNA Polymerization by Cytidine-5′-Phosphoro-2-Aminoimidazolide. Biochemistry 2017, 56 (43), 5739–5747.

(36) Duzdevich, D.; Carr, C. E.; Szostak, J. W. Deep Sequencing of Non-Enzymatic RNA Primer Extension. Nucleic Acids Research 2020, 48 (12), e70–e70.

(37) Chou, F.-C.; Kladwang, W.; Kappel, K.; Das, R. Blind Tests of RNA Nearest-Neighbor Energy Prediction. Proceedings of the National Academy of Sciences 2016, 113 (30), 8430–8435.

(38) Ding, D.; Fang, Z.; Kim, S. C.; O’Flaherty, D. K.; Jia, X.; Stone, T. B.; Zhou, L.; Szostak, J. W. Unusual Base Pair between Two 2-Thiouridines and Its Implication for Nonenzymatic RNA Copying. J. Am. Chem. Soc. 2024, 146 (6), 3861–3871.

(39) Sowers, L. C.; Fazakerley, G. V.; Eritja, R.; Kaplan, B. E.; Goodman, M. F. Base Pairing and Mutagenesis: Observation of a Protonated Base Pair between 2-Aminopurine and Cytosine in an Oligonucleotide by Proton NMR. Proceedings of the National Academy of Sciences 1986, 83 (15), 5434–5438.

(40) Ma, L.; Kartik, S.; Liu, B.; Liu, J. From General Base to General Acid Catalysis in a Sodium-Specific DNAzyme by a Guanine-to-Adenine Mutation. Nucleic Acids Research 2019, 47 (15), 8154–8162.

(41) Handa, S.; Reyna, A.; Wiryaman, T.; Ghosh, P. Determinants of Adenine-Mutagenesis in Diversity-Generating Retroelements. Nucleic Acids Research 2021, 49 (2), 1033–1045.

(42) Rajamani, S.; Ichida, J. K.; Antal, T.; Treco, D. A.; Leu, K.; Nowak, M. A.; Szostak, J. W.; Chen, I. A. Effect of Stalling after Mismatches on the Error Catastrophe in Nonenzymatic Nucleic Acid Replication. J. Am. Chem. Soc. 2010, 132 (16), 5880–5885.

(43) Szostak, J. W. The Narrow Road to the Deep Past: In Search of the Chemistry of the Origin of Life. Angewandte Chemie International Edition 2017, 56 (37), 11037–11043.

(44) Zhang, S. J.; Duzdevich, D.; Ding, D.; Szostak, J. W. Freeze-Thaw Cycles Enable a Prebiotically Plausible and Continuous Pathway from Nucleotide Activation to Nonenzymatic RNA Copying. Proceedings of the National Academy of Sciences 2022, 119 (17), e2116429119.

(45) Aitken, H. R. M.; Wright, T. H.; Radakovic, A.; Szostak, J. W. Small-Molecule Organocatalysis Facilitates In Situ Nucleotide Activation and RNA Copying. J. Am. Chem. Soc. 2023, 145 (29), 16142–16149.

(46) Strobel, S. A.; Cech, T. R.; Usman, N.; Beigelman, L. The 2,6-Diaminopurine Riboside.5-Methylisocytidine Wobble Base Pair: An Isoenergetic Substitution for the Study of G.U Pairs in RNA. Biochemistry 1994, 33 (46), 13824–13835.

