## Supplementary Information for "Diaminopurine in Nonenzymatic RNA Template Copying"

[1] Department of Chemistry and Chemical Biology, Harvard University, 12 Oxford Street, Cambridge, Massachusetts 02138, USA
[2] Department of Molecular Biology and Center for Computational and Integrative Biology, Massachusetts General Hospital, 185 Cambridge Street, Boston, Massachusetts 02114, USA.
[3] Howard Hughes Medical Institute, Department of Chemistry, The University of Chicago, Chicago, Illinois 60637, USA.
[4] Department of Genetics, Harvard Medical School, 77 Avenue Louis Pasteur, Boston, Massachusetts 02115, USA.
[5] Department of Biochemistry and Biophysics, Perelman School of Medicine, University of Pennsylvania, Philadelphia, PA 19104, USA.
[6] Penn Institute for RNA Innovation, University of Pennsylvania, Philadelphia, PA 19104, USA

**1 Materials and Methods**
1.1 General information
1.2 Synthesis and characterization of 2-aminoimidazole activated mononucleotides and 5′-5′ imidazolium bridged dinucleotides
1.3 Synthesis of 2-aminoimidazole activated trinucleotides
1.4 Synthesis of oligonucleotide primers, templates and blocker
1.5 Nonenzymatic primer extension reactions
1.6 Crystallization
1.7 Crystal data collection, structure determination and refinement
1.8 Sequencing
**2 Supplementary Discussion**
RNA Nearest Neighbor Energy Prediction
**3 Supplementary Figures**
**4 Supplementary Tables**

### 1 Materials and Methods

#### 1.1 General information

*Materials.* Reagents and solvents were obtained from Fischer Scientific, Sigma-Aldrich, Alfa Aesar, Acros Organics, Combi-Blocks and were used without any further purification unless otherwise noted. Diaminopurine nucleoside (≥99%) was purchased from ChemImpex. Phosphoramidites and reagents used for solid-phase RNA synthesis were purchased from Glen Research (Sterling, MA) and ChemGenes (Wilmington, MA). Deuterated solvents for NMR were purchased from Cambridge Isotope Laboratories (Tewksbury, MA).

*Nuclear Magnetic Resonance (NMR).* $^{1}H$ and $^{31}P$ NMR spectra were acquired on a Varian Oxford AS-400 NMR spectrometer (400 MHz for $^{1}H$, 162 MHz for $^{31}P$). All spectra were acquired at 25°C. $^{1}H$ NMR was referenced using sodium d6-2,2-dimethyl-2-silapentane-5-sulfonate (DSS) as an internal standard (0 ppm at 25°C).

*Reverse Phase Flash Chromatography.* Purification of 5′-phosphorylated diaminopurine nucleoside, 2-aminoimidazole activated mononucleotide, 5′-5′ imidazolium bridged dinucleotides and 5′-phosphorylated trinucleotides was performed using a prepacked RediSep Rf Gold C18Aq 50 g column from Teledyne Isco (Lincoln, NE).

*Preparatory-scale High Performance Liquid Chromatography (prep-HPLC).* Purification of activated trinucleotide helpers was carried out on an Agilent 1290 HPLC system, equipped with a preparative-scale Agilent ZORBAX Eclipse-XDB C18 column (21.2x250mm, 7 µm particle size).
Low-resolution Mass Spectrometry (LRMS). All samples were diluted to 200 µM in Milli-Q water (18.2 MΩ·cm) with a few drops of acetonitrile immediately prior to analysis. The spectra were obtained by direct injection on an Esquire 6000 mass spectrometer (Bruker Daltonics), operated in the alternating ion mode.

*High-resolution Liquid Chromatography Mass Spectrometry (HR LC-MS).* The samples were separated and analyzed using an Agilent 1200 High-performance Liquid Chromatography (HPLC) coupled to an Agilent 6230 Time-of-flight Mass Spectrometry (TOF MS) equipped with a diode array detector. The samples were separated by IP-RP-HPLC on a 100 mm × 1 mm (length × i.d.) Xbridge C18 column with 3.5 µm particle size (Waters, Milford, MA). The samples were eluted between 2.5 and 15% methanol in 200 mM 1,1,1,3,3,3-hexafluoro-2propanol with 1.25 mM triethylamine at pH 7.0 over 16 minutes with a flow rate of 0.1 mL/min at 50°C. The samples were analyzed in negative mode from 239 m/z to 3200 m/z with a scan rate of 1 spectrum/s.

**1.2 Synthesis and characterization of 2-aminoimidazole activated mononucleotides and 5′-5′ imidazolium bridged dinucleotides**

*Phosphorylation of diaminopurine nucleoside.* The 5′-OH of the diaminopurine nucleoside was phosphorylated using a previously reported procedure.[1] To a pre-chilled mixture of diaminopurine nucleoside (1 equiv.) in $OP(OMe)_3$ (0.1 M with respect to the nucleoside) was added $POCl_3$ (4 equiv.) under vigorous stirring at 0°C. After complete solubilization of the nucleoside, DIPEA (0.5 eq.) was added drop-wise to the stirring reaction. Three additional portions of DIPEA were added (0.5 eq. each) at 20 minute intervals. Once the starting material disappeared, monitored by LRMS, the reaction was quenched using 1 M TEAB (5X volumes, pH 7.5). The product was purified by reverse phase flash chromatography using gradient elution between (A) aqueous 2 mM TEAB (pH 7.5) and (B) acetonitrile. The product was eluted between 0% and 15% B over 13 CVs with a flow rate of 40 mL/min. Fractions containing product were collected and lyophilized. The diaminopurine nucleotide was used for downstream activation without further purification.

*Synthesis and characterization of 2-aminoimidazole activated mononucleotides (*N).* The synthesis of *A, *U, *C, *G and *D follows a previously reported procedure.[2] The detailed characterization (NMR and HR-MS) of *A, *U, *C, *G can be found in reference 2 and that of *D is listed below.

ribo-diaminopurine-5′-phosphoro-(2-aminoimidazolide) **(*D)**

$^1$H NMR (400 MHz, $D_2O$) δ 7.98 (s, 1H), 6.67 – 6.61 (m, 1H), 6.48 – 6.41 (m, 1H), 5.89 (d, *J* = 5.7 Hz, 1H), 4.73 (t, *J* = 5.4 Hz, 1H), 4.41 – 4.36 (m, 1H), 4.29 – 4.24 (m, 1H), 4.12 – 4.03 (m, 2H). Peaks corresponding to residual TEAB observed at 3.20 and 1.27 ppm.

**$^{31}$P NMR** (162 MHz, $D_2O$) δ -11.44.

**HRMS** (Q-TOF) m/z: $[M-H]^-$ calculated for $C_{13}H_{17}N_9O_6P$ 426.1039; found: 426.1097.

*Synthesis and characterization of 5′-5′ imidazolium-bridged dinucleotides.* The synthesis of A*A and D*D follows a previously reported procedure.[3] The detailed characterization (NMR and HR-MS) of A*A can be found in reference 3 and that of D*D is listed below.

1,3-di-(diaminopurine-5′phosphoryl)-2-aminoimidazolium **(D*D)**

**$^1$H NMR** (400 MHz, $D_2O$) δ 7.81 (s, 2H), 6.65 (s, 2H), 5.76 (d, *J* = 5.0 Hz, 2H), 4.59 (t, *J* = 5.2 Hz, 1H), 4.37 (t, *J* = 4.8 Hz, 2H), 4.15 – 4.01 (m, 4H), 4.00 – 3.85 (m, 2H). Peaks corresponding to residual TEAB observed at 3.20 and 1.27 ppm.

**$^{31}$P NMR** (162 MHz, $D_2O$) δ -12.77.

**HRMS** (Q-TOF) m/z: $[M-H]^-$ calculated for $C_{23}H_{30}N_{15}O_{12}P_2$ 770.1679; found: 770.1747.

#### 1.3 Synthesis of 2-aminoimidazole activated trinucleotides

The 5′-phosphorylated trinucleotides (GAC and AGG) were prepared by solid-phase synthesis on a MerMade 6 DNA/RNA synthesizer. The trinucleotides were subsequently deprotected and purified by reverse phase flash chromatography on a 50 g C18Aq column over 12 CVs of 0-10% acetonitrile in 2 mM TEAB buffer (pH 8.0). The fractions corresponding to the 5′-phosphorylated trinucleotides, as monitored by LRMS, were collected and lyophilized to complete dryness. Trinucleotides (1 equiv.), 2-aminoimidazole hydrochloride (2AI·HCl, 40 equiv.), and triphenylphosphine (TPP, 40 equiv.) were dissolved in dry dimethylsulfoxide (DMSO) with triethylamine (TEA, 400 equiv.). 2,2′-dipyridyldisulfide (DPDS, 40 equiv.) was added to initiate the activation and the solution was incubated at room temperature for 6 hours. The solution was then precipitated and purified on prep-HPLC over 12 CVs of 2-8% acetonitrile in 20 mM TEAB buffer (pH 8.0). The fractions corresponding to 2AI activated trinucleotides (*GAC and *AGG), as monitored by LRMS, were collected and pH adjusted to 10 using 10 M NaOH. The activated trinucleotides were then aliquoted into 50 nmol fractions and lyophilized. The detailed characterizations (HR-MS) are included in the supplementary data.

#### 1.4 Synthesis of oligonucleotide primers, templates and blocker

The oligonucleotides made of canonical nucleobases were purchased from Integrated DNA Technologies (Coralville, IA). The oligonucleotides containing D were prepared on an Expedite 8909 DNA/RNA synthesizer. The oligonucleotides were cleaved from the solid support and deprotected with AMA (ammonium hydroxide/40% aqueous methylamine 1:1 v/v) at 65°C for 20 minutes. The mixtures were lyophilized, 2′-deprotected by removal of the 2′-TBDMS groups and purified using Glen-pak RNA cartridges (Glen research, Sterling, MA). The oligonucleotides were then lyophilized and resuspended in 7M urea to be further purified by 20% (19:1) polyacrylamide gel electrophoresis (PAGE, National Diagnostics, Atlanta, GA) and Sep-pak C18 plus short cartridges (Waters, Milford, MA).

#### 1.5 Nonenzymatic primer extension reactions

*Michaelis-Menten kinetics.* The annealing solutions consisting of the primer/template/blocker complexes were prepared at 5X final concentration: 7.5 µM primer, 12.5 µM template, 17.5 µM blocker, 50 mM Tris-Cl pH 8.0, 50 mM NaCl, and 1 mM EDTA. The solution was heated at 85°C for 30 s and then slowly cooled to 25°C at a rate of 0.1°C/s in a thermal cycler machine. The annealed solution was diluted to yield the resulting final concentrations: 1.5 µM primer, 2.5 µM template, 3.5 µM blocker, 200 mM Tris-Cl pH 8.0, and 100 mM $MgCl_2$. Stock solutions of bridged dinucleotides (A*A or D*D), freshly prepared at 2X titrating concentrations, were added to the annealed primer/template/blocker solution to initiate the templated primer extension reactions. At each time point, 0.5 µL of reaction sample was added to 25 µL quenching buffer containing 25 mM EDTA, 1X TBE, and 4 µM of an DNA sequence complementary to the template in formamide. The sequences of oligonucleotides can be found in Table S1.

*Primer extension reactions with activated mononucleotide and downstream activated trinucleotide helper.* The annealing solution consisting of primer/template complexes were prepared at 5X final concentration: 7.5 µM primer, 12.5 µM template, 50 mM Tris-Cl pH 8.0, 50 mM NaCl, and 1 mM EDTA. The solution was annealed as previously described and diluted to yield the resulting final concentrations: 1.5 µM primer, 2.5 µM template, 200 mM Tris-Cl pH 8.0, and 100 mM $MgCl_2$. For initiating the reactions, activated mononucleotides (*A, *D, *U, *C, *G) and activated trinucleotides (*GAC or *AGG) were freshly prepared and added to the annealed primer/template solution to yield the final concentrations of 20 mM activated mononucleotides and 0.5 mM activated trinucleotides. At each time point, the reaction was quenched as previously described. The sequences of oligonucleotides can be found in Table S4.

*Stalling effect of the D:C mismatch.* The reaction conditions of the stalling experiments are as described in the primer extension reactions with activated mononucleotide and downstream activated trinucleotide except that the reactions were initiated by adding bridged dinucleotides (G*G) to a final concentration of 10 mM. The sequences of oligonucleotides can be found in Table S5.

#### 1.6 Crystallization

0.33 mM self-complementary RNA sequences in nuclease-free water (Invitrogen, Waltham, MA) were heated to 90°C for 2 minutes and then slowly cooled to room temperature. Crystal Screen

HT, Index HT, Natrix HT (Hampton Research, Aliso Viejo, CA) and Nuc-Pro HTS (Jena Bioscience, Jena, Germany) were used to screen crystallization conditions at 20°C using the sitting-drop vapor diffusion method. An NT8 robotic system and Rock Imager (Formulatrix, Waltham, MA) were used for crystallization screening and monitoring the crystallization process. The sequences of self-complementary RNA duplexes are listed in Table S6 and the optimal crystallization conditions are listed in the Table S7.

**1.7 Crystal data collection, structure determination and refinement**

Diffraction data were collected under a liquid nitrogen stream at 99 K at a wavelength of 1.038413 Å or 1.033216 Å on Beamline 201 at the Advanced Light Source in the Lawrence Berkeley National Laboratory (USA). The crystals were exposed for 0.25 s per image with a 0.25° oscillation angle on Beamline 201. The distances between detector and the crystal were set to 200-300 mm. The data were processed by HKL2000[4] or XDS. The structures were solved by molecular replacement by PHASER[5] using the structure of 3ND4 as the searching model[6]. All structures were refined by Refmac5 in CCP4i[7] or Phenix[8]. After several cycles of refinement, some water molecules were added in Coot[9]. Data collection, phasing, and refinement statistics of the determined structures are listed in Supplementary Table S8 and S9.

**1.8 Sequencing**

*RNA sample preparation for Illumina Sequencing.* The procedure for RNA sample preparation for sequencing is adapted from the reported protocol.[10] Table S10 lists all sequences used for the sequencing experiments. The annealing solution consisted of the self-complementary hairpin constructs were prepared at 5X final concentration: 5 µM hairpin construct (oligo 6N for AUCG system, and oligo 6D for DUCG system, Table S10), 6 µM blocker, 50 mM Tris-Cl pH 8.0, 1 mM EDTA, 50 mM NaCl. The solution was annealed as previously described and diluted to yield the resulting final concentrations: 1 µM hairpin construct, 1.2 µM blocker, 200 mM Tris-Cl pH 8.0, and indicated amount of $MgCl_2$ (10 mM or 100 mM). Bridged dinucleotides (N*N) mixtures (for AUCG system, N=A, U, C, G; for DUCG system, N=D, U, C, G) were obtained by equilibrating freshly prepared bridged homo-dinucleotides at room temperature for 2 hours (for AUCG system, equilibration of A*A, U*U, C*C and G*G; for DUCG system, equilibration of D*D, U*U, C*C and G*G).[11] N*N mixtures were added to the solution at a 10 mM or 20 mM final concentration to initiate the reactions. The final volume of the reactions was 30 μL. The reactions were incubated at room temperature for 24 hours and quenched by adding 20 μL 0.5 M EDTA.

The reactions were subsequently desalted using the Nu-Clean D25 centrifuge spin columns (IBI Scientific, Dubuque, IA). The NPOM caged bases were uncaged under 385 nm UV for 45 minutes. The samples were purified by 20% (19:1) polyacrylamide gel electrophoresis (PAGE) and recovered by ZR small-RNA PAGE Recovery Kit (Zymo Research, Tustin, CA). The RT Handle (template for the reverse transcription primer) was ligated using T4 RNA Ligase 2, truncated KQ (New England Biolabs, Ipswich, MA). The samples were treated with Proteinase K (New England Biolabs), extracted with Phenol/Chloroform/Isoamyl Alcohol (25:24:1) (Fischer Scientific, Hampton, NH), and concentrated using the Oligo Clean & Concentrator Kits (Zymo Research). The RT Primer was annealed to the purified sample and the RNA oligo was reverse transcribed with ProtoScript II Reverse Transcriptase (New England Biolabs). The RNA in the resulting mixtures were hydrolyzed using 1M NaOH. The reactions were purified using the Oligo Clean & Concentrator Kits and eluted in 20 μL TE buffer, pH 7.0. 10 uL of the recovered cDNA solution was added to the 40 uL reaction mixtures containing 0.5 μL of Q5 HotStart High-Fidelity DNA Polymerase PCR reaction, 1 μL of NEBNext SR Primer for Illumina and 1 μL NEBNextIndex Index Primer for Illumina (New England Biolabs). The reactions were run for 6 cycles with a 15 s 62°C extension step.

The PCR products were purified by agarose gel electrophoresis with 3:1 mixture of Metaphor agarose (Lonza, Basel, Switzerland) and agarose (Bio-Rad, Hercules, CA). The products were recovered by Quantum Prep Freeze 'N Squeeze DNA Gel Extraction Spin Columns (Bio-Rad) and subsequently by Agencourt AMPure XP magnetic beads (Beckman Coulter, Brea, CA). The size and the concentration of the samples were measured by TapeStation (4200 TapeStation System, Agilent) and concentration was verified by qPCR (CFX384 Real-Time System, Bio-Rad). The samples were pooled with 30% PhiX spiked in (PhiX Control v3, Illumina, San Diego, CA) and submitted for paired-end sequencing by MiSeq (MiSeq Reagent Kit v3, 150 cycles, Illumina).

*Sequencing data analysis.* The raw sequencing files were analyzed by the NERPE-Seq code written in MATLAB[10] to generate the output data. The output data were further processed by Excel and Python, and represented by Prism (v9.5.1) to generate the yield and fidelity, product base and bridged dinucleotides distributions, template:product pair distribution, and mismatch pattern plots.

The raw sequencing files, processing worksheets and code can be found in the OSF.io repository: https://osf.io/zscy8/.

### 2 Supplementary Discussion

#### RNA Nearest Neighbor Energy Prediction

Based on the equation $K_d = e^{\frac{\Delta G}{RT}}$, then $\Delta\Delta G = \ln \frac{{K_d}^{DD}}{{K_d}^{AA}} \times RT$. $K_d$ can be approximated using $K_m$ by assuming that the on and off rates of the imidazolium-bridged dinucleotide substrate are fast relative to the chemical step.[3] The equation then becomes $\Delta\Delta G = \ln \frac{{K_m}^{DD}}{{K_m}^{AA}} \times RT$. Inserting the values of ${K_m}^{AA}$ and ${K_m}^{DD}$ from Figure 2C, we computed $\Delta\Delta G°_{25} = -1.757\ \text{kcal/mol}$.

According to the nearest-neighbor (NN) model[12], for the RNA duplex construct used in Figure 2 (Table S1), the Gibbs free energies of AA and DD systems are calculated as:

$\Delta G°_{37}{}^{AA} = \Delta G°_{37,\text{intermolecular initiation}} + \Delta G°_{37,GA/CU} + \Delta G°_{37,AA/UU} + \Delta G°_{37,\text{symmetry}} + \Delta G°_{37,\text{AU end penalty}}$ and

$\Delta G°_{37}{}^{DD} = \Delta G°_{37,\text{intermolecular initiation}} + \Delta G°_{37,GD/CU} + \Delta G°_{37,DD/UU} + \Delta G°_{37,\text{symmetry}} + \Delta G°_{37,\text{DU end penalty}}$

, with coaxial stacking energies approximated using the NN values for contiguous base pairs. Assuming $\Delta G°_{37,\text{intermolecular initiation}}$ and $\Delta G°_{37,\text{symmetry}}$ are constant in both equations, the change in Gibbs free energy becomes:

$$\Delta\Delta G°_{37} = \Delta G°_{37}{}^{DD} - \Delta G°_{37}{}^{AA}$$
$$= (\Delta G°_{37,GD/CU} - \Delta G°_{37,GA/CU}) + (\Delta G°_{37,DD/UU} - \Delta G°_{37,AA/UU}) + (\Delta G°_{37,\text{DU end penalty}} - \Delta G°_{37,\text{AU end penalty}}).$$

We chose to use the RECCES-Rosetta predicted NN parameters[13] because the experimentally reported NN parameters and molecular dynamics/quantum mechanics (MD/QM) predictions have missing values (Table S2). The above equation simplifies to:

$$\Delta\Delta G°_{37} = \Delta G°_{37}{}^{DD} - \Delta G°_{37}{}^{AA}$$
$$= \big((-3.10 \pm 0.17) - (-2.13 \pm 0.09)\big) + \big((-3.46 \pm 0.18) - (-1.13 \pm 0.17)\big) + \big((-0.02 \pm 0.19) - (0.76 \pm 0.15)\big)$$
$$= -4.08 \pm 0.40\ \text{kcal/mol}$$

Our calculated change in Gibbs free energy is less negative than that derived from the Rosetta predicted NN parameters. In addition to Rosetta predictions, experimentally reported NN parameters and molecular dynamics/quantum mechanics (MD/QM) predictions indicate slightly different ΔG values. It is important to note that the trend across all three categories is consistent: the ΔG associated with D is in general smaller than that with A (Table S2). Consequently, the negative change in Gibbs free energy predicted by the NN model indicates that substituting A with D can increase the thermodynamic stability of RNA duplexes, a conclusion that is in alignment with the trends observed in our experimentally measured value.

### References


(1) Yoshikawa, M.; Kato, T.; Takenishi, T. A Novel Method for Phosphorylation of Nucleosides to 5′-Nucleotides. *Tetrahedron Letters* **1967**, *8* (50), 5065–5068.
(2) Li, L.; Prywes, N.; Tam, C. P.; O'Flaherty, D. K.; Lelyveld, V. S.; Izgu, E. C.; Pal, A.; Szostak, J. W. Enhanced Nonenzymatic RNA Copying with 2-Aminoimidazole Activated Nucleotides. *J. Am. Chem. Soc.* **2017**, *139* (5), 1810–1813.
(3) Ding, D.; Zhou, L.; Giurgiu, C.; Szostak, J. W. Kinetic Explanations for the Sequence Biases Observed in the Nonenzymatic Copying of RNA Templates. *Nucleic Acids Res* **2022**, *50* (1), 35–45.
(4) Otwinowski, Z.; Minor, W. Processing of X-Ray Diffraction Data Collected in Oscillation Mode. In *Methods in Enzymology*; Macromolecular Crystallography Part A; Elsevier, 1997; Vol. 276, pp 307–326.
(5) McCoy, A. J.; Grosse-Kunstleve, R. W.; Adams, P. D.; Winn, M. D.; Storoni, L. C.; Read, R. J. Phaser Crystallographic Software. *J Appl Crystallogr* **2007**, *40* (4), 658–674.
(6) Mooers, B. H. M.; Singh, A. The Crystal Structure of an Oligo(U):Pre-mRNA Duplex from a Trypanosome RNA Editing Substrate. *RNA* **2011**, *17* (10), 1870–1883.
(7) Murshudov, G. N.; Vagin, A. A.; Dodson, E. J. Refinement of Macromolecular Structures by the Maximum-Likelihood Method. *Acta Crystallogr D Biol Crystallogr* **1997**, *53* (3), 240–255.
(8) Liebschner, D.; Afonine, P. V.; Baker, M. L.; Bunkóczi, G.; Chen, V. B.; Croll, T. I.; Hintze, B.; Hung, L. W.; Jain, S.; McCoy, A. J.; Moriarty, N. W.; Oeffner, R. D.; Poon, B. K.; Prisant, M. G.; Read, R. J.; Richardson, J. S.; Richardson, D. C.; Sammito, M. D.; Sobolev, O. V.; Stockwell, D. H.; Terwilliger, T. C.; Urzhumtsev, A. G.; Videau, L. L.; Williams, C. J.; Adams, P. D. Macromolecular Structure Determination Using X-Rays, Neutrons and Electrons: Recent Developments in Phenix. *Acta Crystallogr D Struct Biol* **2019**, *75* (10), 861–877.
(9) Emsley, P.; Lohkamp, B.; Scott, W. G.; Cowtan, K. Features and Development of Coot. *Acta Crystallogr D Biol Crystallogr* **2010**, *66* (4), 486–501.
(10) Duzdevich, D.; Carr, C. E.; Szostak, J. W. Deep Sequencing of Non-Enzymatic RNA Primer Extension. *Nucleic Acids Research* **2020**, *48* (12), e70–e70.
(11) Ding, D.; Zhou, L.; Mittal, S.; Szostak, J. W. Experimental Tests of the Virtual Circular Genome Model for Nonenzymatic RNA Replication. *J. Am. Chem. Soc.* **2023**, *145* (13), 7504–7515.
(12) Turner, D. H.; Mathews, D. H. NNDB: The Nearest Neighbor Parameter Database for Predicting Stability of Nucleic Acid Secondary Structure. *Nucleic Acids Research* **2010**, *38* (suppl_1), D280–D282.
(13) Chou, F.-C.; Kladwang, W.; Kappel, K.; Das, R. Blind Tests of RNA Nearest-Neighbor Energy Prediction. *Proceedings of the National Academy of Sciences* **2016**, *113* (30), 8430–8435.
(14) Xia, T.; SantaLucia, J. Jr.; Burkard, M. E.; Kierzek, R.; Schroeder, S. J.; Jiao, X.; Cox, C.; Turner, D. H. Thermodynamic Parameters for an Expanded Nearest-Neighbor Model for Formation of RNA Duplexes with Watson−Crick Base Pairs. *Biochemistry* **1998**, *37* (42), 14719–14735.
(15) Chen, J. L.; Dishler, A. L.; Kennedy, S. D.; Yildirim, I.; Liu, B.; Turner, D. H.; Serra, M. J. Testing the Nearest Neighbor Model for Canonical RNA Base Pairs: Revision of GU Parameters. *Biochemistry* **2012**, *51* (16), 3508–3522.

(16) Hopfinger, M. C.; Kirkpatrick, C. C.; Znosko, B. M. Predictions and Analyses of RNA Nearest Neighbor Parameters for Modified Nucleotides. *Nucleic Acids Research* **2020**, *48* (16), 8901–8913.

### 3 Supplementary Figures

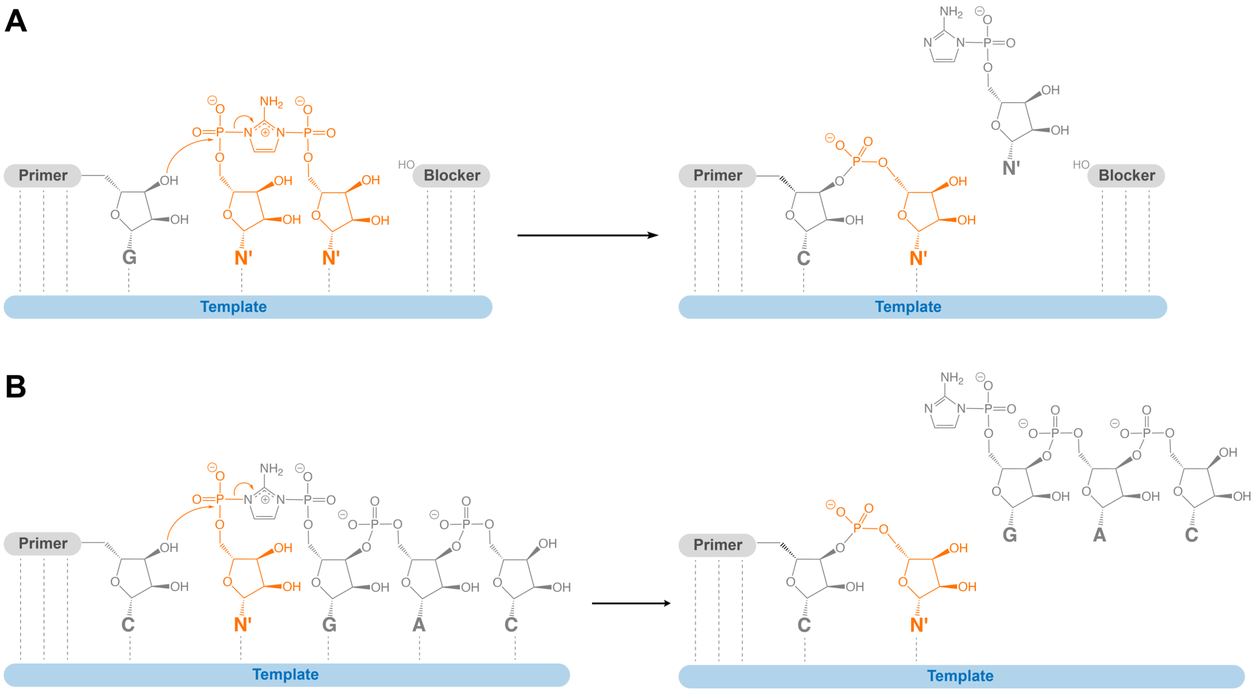


**Figure S1.** Mechanism of nonenzymatic primer extension reactions through (A) bridged dinucleotide intermediate inside a primer-template-blocker system and (B) monomer-bridged-trimer intermediate.

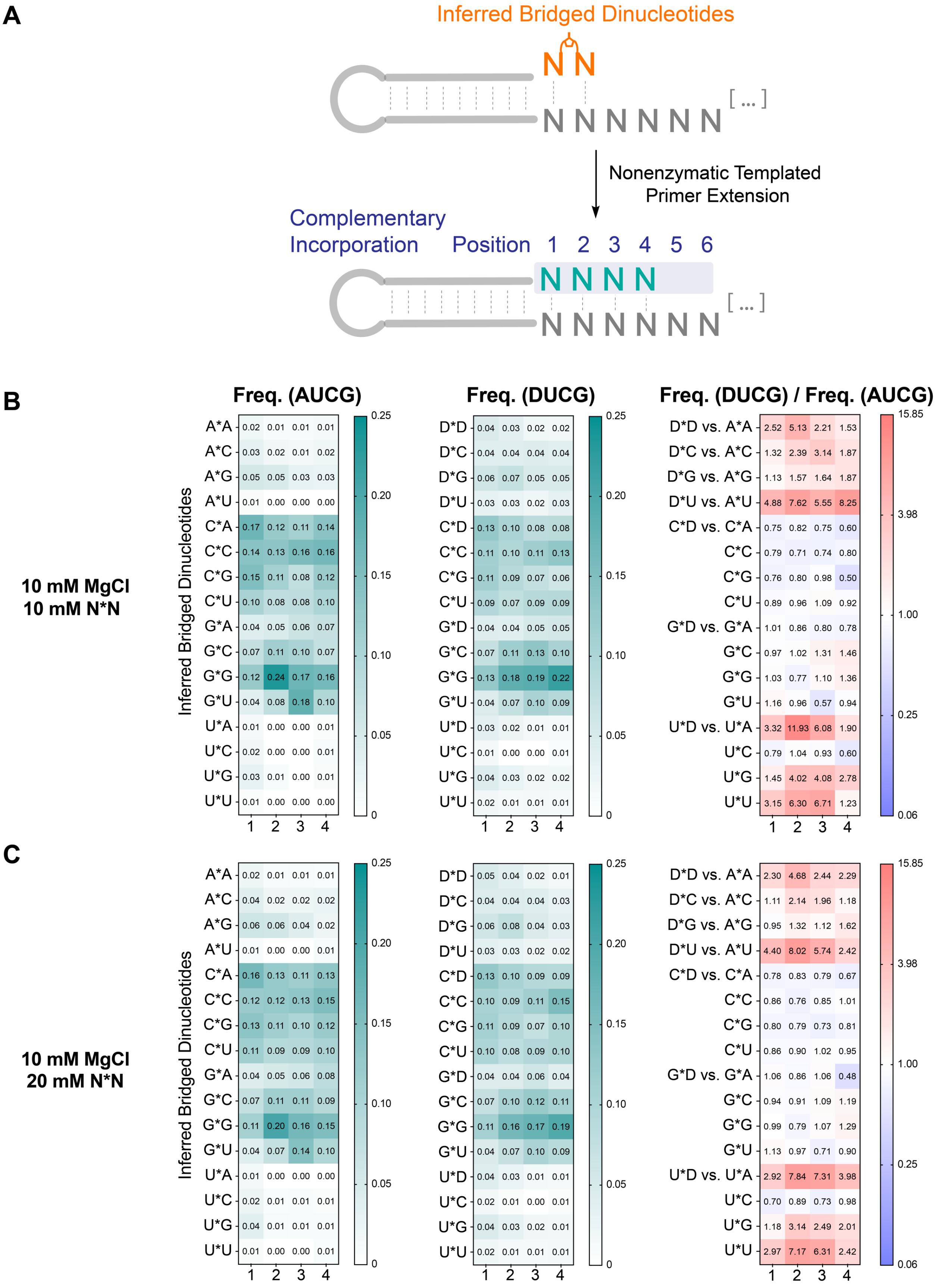

A
Inferred Bridged Dinucleotides
Nonenzymatic Templated Primer Extension
Complementary Incorporation
Position 1 2 3 4 5 6
B
Freq. (AUCG)
Freq. (DUCG)
Freq. (DUCG) / Freq. (AUCG)
10 mM MgCl
10 mM N*N
Inferred Bridged Dinucleotides
C
10 mM MgCl
20 mM N*N
Inferred Bridged Dinucleotides

**D**

**100 mM MgCl**
**20 mM N*N**

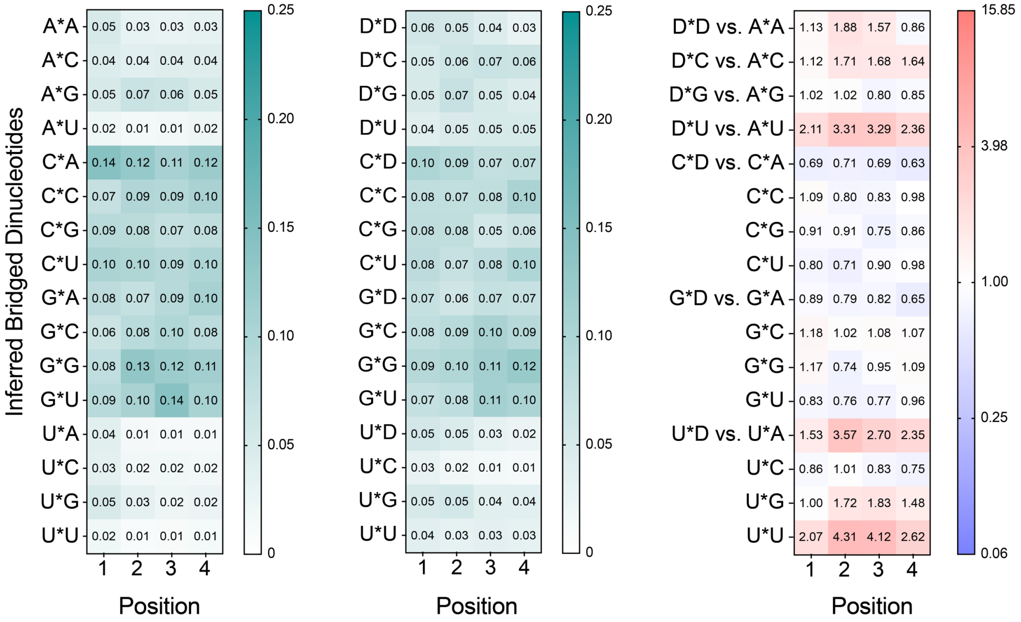


**Figure S2.** DUCG system decreases biases in complementary product incorporation by enriching the distribution of D*N and U*N bridged dinucleotides. (A) Schematic representation of the bridged dinucleotides' distribution. The exact sequences of the bridged dinucleotides are inferred from the template. (B-D) Position-dependent frequency of bridged dinucleotides in the AUCG & DUCG systems and the frequency ratio between AUCG and DUCG. Heatmaps are generated for the following reaction conditions: (B) 10 mM $MgCl_2$ and 10 mM N*N (C) 10 mM $MgCl_2$ and 20 mM N*N (D) 100 mM $MgCl_2$ and 20 mM N*N. For the frequency ratio heatmap, red represents greater frequency in the DUCG system whereas blue represents greater frequency in the AUCG system.

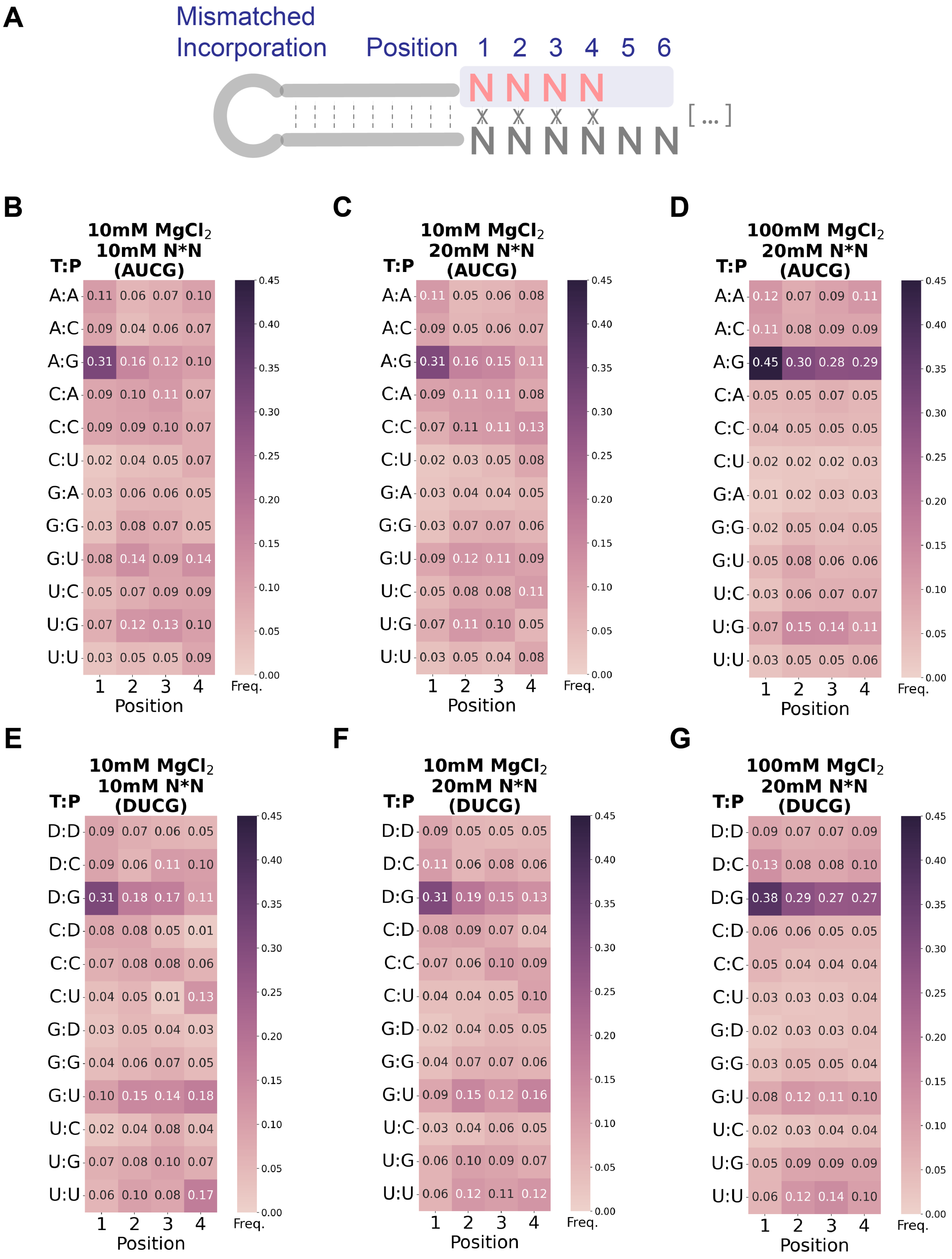


**Figure S3.** Mismatch heatmaps show the position-dependent distribution of mismatched pairs (template: product) among the mismatched incorporation. (A) Schematic representation of the mismatched incorporation. At each position, the frequencies of all 12 possible T:P mismatch pairs are normalized to 1. (B-G) Heatmaps are generated for the following reaction conditions: 10 mM $MgCl_2$ and 10 mM N*N in (B) AUCG and (E) DUCG systems. 10 mM $MgCl_2$ and 20 mM N*N in (C) AUCG and (F) DUCG systems. 100 mM $MgCl_2$ and 20 mM N*N in (D) AUCG and (G) DUCG systems.

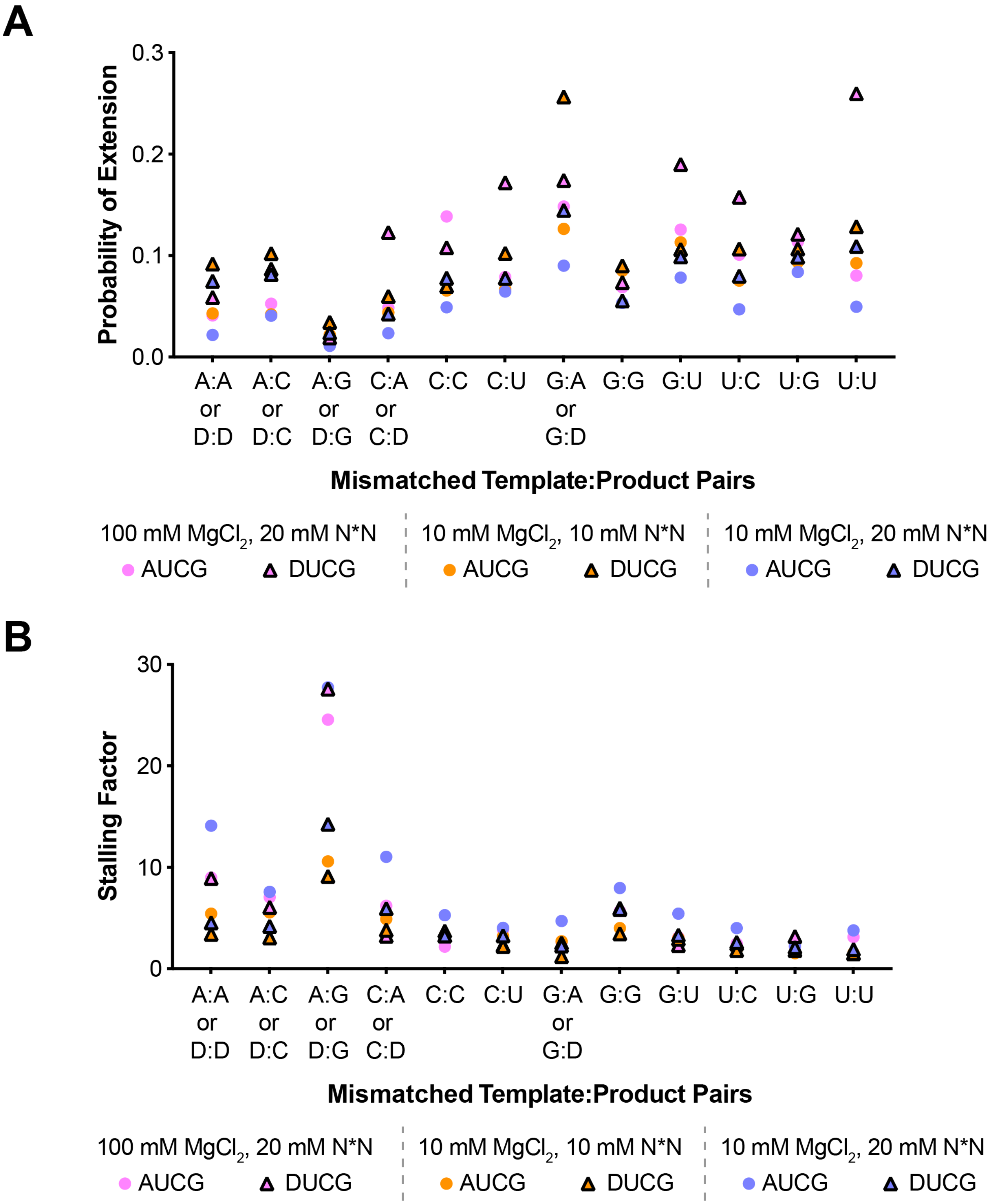


**Figure S4.** The effect of mismatches at position 1 on subsequent primer extension. (A) Extension probability over each mismatch at position 1. (B) Stalling factor for each type of mismatch at position 1, defined as the ratio of the extension probability after a complementary pair compared to that after a mismatched pair.

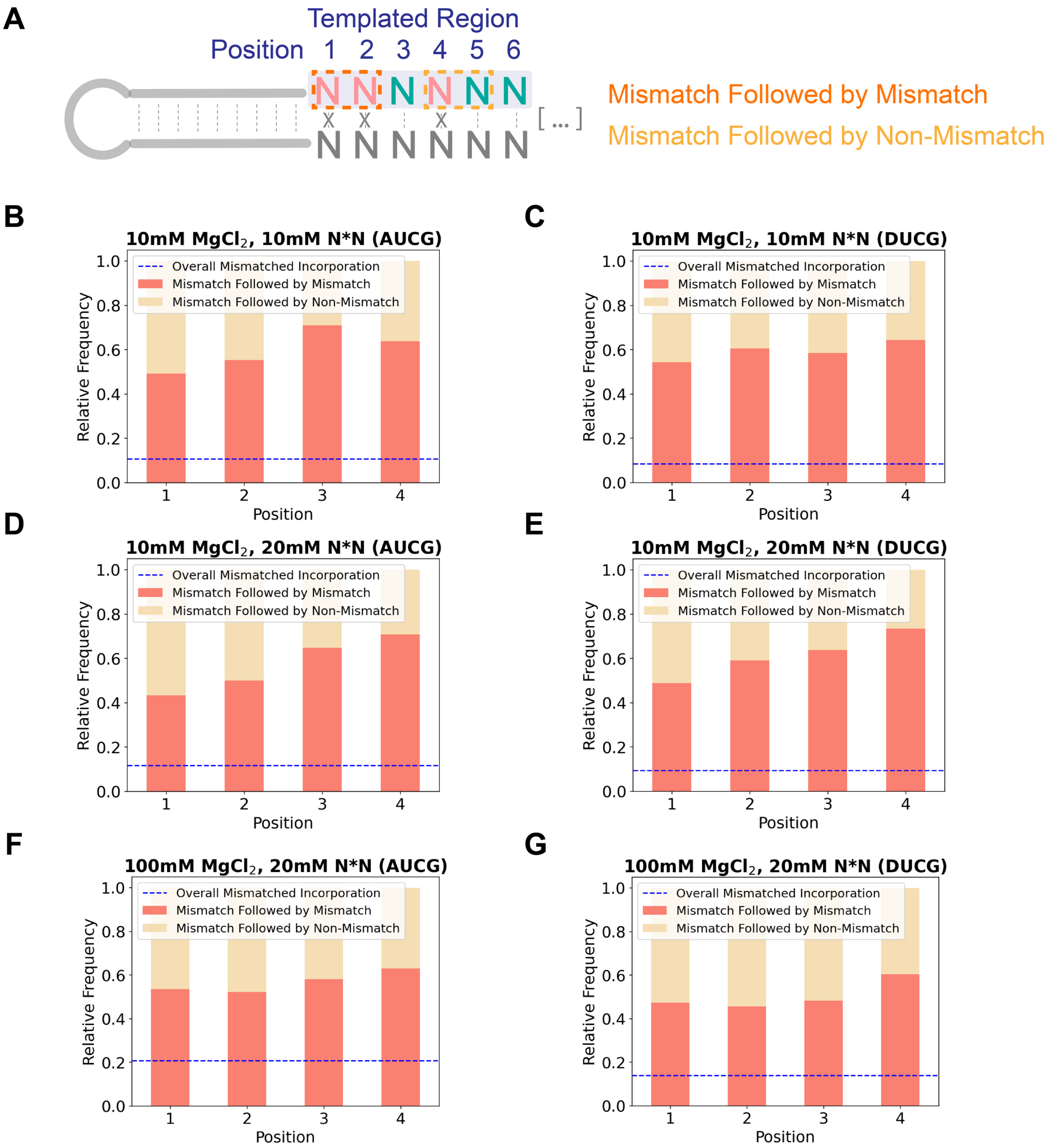


**Figure S5.** Extending mismatch stacked barplots show the position-dependent relative frequency of mismatch followed by mismatch and non-mismatch at position 1-4 in AUCG and DUCG systems. (A) Schematic representation of mismatch followed by mismatch and non-mismatch. At each position, the frequencies of mismatch followed by mismatch and mismatch followed by non-mismatch are normalized to 1. (B-G) Stacked barplots are generated for the following reaction conditions: 10 mM $MgCl_2$ and 10 mM N*N in (B) AUCG and (C) DUCG systems. 10 mM $MgCl_2$ and 20 mM N*N in (D) AUCG and (E) DUCG systems. 100 mM $MgCl_2$ and 20 mM N*N in (F) AUCG and (G) DUCG systems.

### 4 Supplementary Tables

**Table S1. Sequences of the primer, template, blocker, and complementary oligonucleotide used in the Michaelis-Menten reactions.**

| Name | Role | Source | Type | Sequence (5′→ 3′) |
|---|---|---|---|---|
| DL-30 | Primer | IDT | RNA | /FAM/AGU GAG UAA CGG |
| LA-122 | Template | IDT | RNA | AU AUC UGA CAU CUU CCG UUA CUC ACU |
| LA-111 | Blocker | IDT | RNA | G AUG UCA GAU AU |
| dCLA-122 | Complementary strand | IDT | DNA | AGT GAG TAA CGG AAG ATG TCA GAT AT |

**Table S2. RECCES-Rosetta predictions[13], experimentally reported NN parameters[14,15], and MD/QM predictions[16]. All values are reported in kcal/mol.**

| NN Sequence | $\Delta G°_{37,\text{ RECCES-Rosetta}}$ | $\Delta G°_{37,\text{ exp}}$ | $\Delta G°_{37,\text{ MD/QM}}$ |
|---|---|---|---|
| 5′ GA 3′<br>3′ CU 5′ | -2.13 ± 0.09 | -2.35 ± 0.06 | -2.18 ± 0.07 |
| 5′ GD 3′<br>3′ CU 5′ | -3.10 ± 0.17 | -3.10 ± 0.21 | -2.37 ± 0.09 |
| 5′ AA 3′<br>3′ UU 5′ | -1.13 ± 0.17 | -0.93 ± 0.03 | -1.02 ± 0.12 |
| 5′ DD 3′<br>3′ UU 5′ | -3.46 ± 0.18 | N/A | N/A |
| 5′ AU 3′<br>3′ UA 5′ | -0.91 ± 0.21 | -1.10 ± 0.08 | -1.11 ± 0.13 |
| 5′ DU 3′<br>3′ UA 5′ | -1.85 ± 0.19 | N/A | -1.41 ± 0.08 |
| Terminal A:U | 0.76 ± 0.15 | 0.45 ± 0.04 | N/A |
| Terminal D:U | -0.02 ± 0.19 | N/A | N/A |

**Table S3. Stacking interaction energy differences predicted by the MD/QM calculations[16]. NN pairs are within the 5′$X_1NY_1$/3′$X_2UY_2$ trimers (N = A or D). The summation of stacking interaction energies is from 4 pairs of stacking interactions (2 intra-strand and 2 inter-strand stackings). All values are reported in kcal/mol.**

| NN type [1] | NN-1 | NN-2 | Σ Stacking Energies (NN-2)<br>- Σ Stacking Energies (NN-1) |
|---|---|---|---|
| $X_1$ N<br>$X_2$U | AA<br>UU | AD<br>UU | -0.95 ± 0.20 |
| $X_1$ N<br>$X_2$U | UA<br>AU | UD<br>AU | -0.59 ± 0.31 |
| $NY_1$<br>$UY_2$ | AA<br>UU | DA<br>UU | -0.57 ± 0.10 |
| $NY_1$<br>$UY_2$ | AU<br>UA | DU<br>UA | -0.50 ± 0.20 |

[1] N in NN type stands for A or D.

**Table S4. Sequences of the primer, templates, and complementary oligonucleotides used in the primer extension reactions with activated mononucleotide and downstream activated trinucleotide.**

| Name | Role | Source | Type | Sequence (5′→ 3′) [1] |
|---|---|---|---|---|
| DU-1 | Primer | IDT | RNA | /FAM/AGU GAG UAA CGC |
| DU-2 | Template | IDT | RNA | GUC C GCG UUA CUC ACU |
| CDU-2 | Complementary strand | IDT | RNA | AGU GAG UAA CGC G GAC |
| DU-3-0 | Template | IDT | RNA | GUC A GCG UUA CUC ACU |
| DU-3 | Template | In-house | RNA | GUC D GCG UUA CUC ACU |
| CDU-3 | Complementary strand | IDT | RNA | AGU GAG UAA CGC U GAC |
| DU-4 | Template | IDT | RNA | GUC G GCG UUA CUC ACU |
| CDU-4 | Complementary strand | IDT | RNA | AGU GAG UAA CGC C GAC |
| DU-5 | Template | IDT | RNA | GUC U GCG UUA CUC ACU |
| CDU-5 | Complementary strand | IDT | RNA | AGU GAG UAA CGC A GAC |
| DU-12 [2] | Template | IDT | RNA | CCUC GCG UUA CUC ACU |
| CDU-12 [2] | Complementary strand | IDT | RNA | AGU GAG UAA CGC GAGG |

[1] Underlined bases vary in each set of sequences (template and complementary strand) while the other bases of the oligonucleotides hold the same.
[2] DU-6 and CDU-6 are used with activated trimer sequence of *AGG instead of *GAC in all other sequences.

**Table S5. Sequences of the primers, templates, and complementary oligonucleotides used in the stalling effect experiments.**

| Name | Role | Source | Type | Sequence (5′→ 3′) [1] |
|---|---|---|---|---|
| DU-6 | Primer | In-house | RNA | /FAM/ AGU GAG UAA CGC <u>D</u> |
| DU-7 | Primer | IDT | RNA | /FAM/ AGU GAG UAA CGC <u>G</u> |
| DU-10 | Template | IDT | RNA | AA CCC<u>C</u> GCG UUA CUC ACU |
| CDU-10 | Complementary strand | IDT | RNA | AGU GAG UAA CGC <u>G</u>GGG UU |
| DU-8 | Primer | IDT | RNA | /FAM/ AGU GAG UAA CGC C |
| DU-9 | Primer | IDT | RNA | /FAM/ AGU GAG UAA CGC U |
| DU-11 | Template | In-house | RNA | AA CCC<u>D</u> GCG UUA CUC ACU |
| CDU-11 | Complementary strand | IDT | RNA | AGU GAG UAA CGC <u>U</u>GGG UU |

[1] Underlined bases vary in each set of sequences (primer, template and complementary strand) while the other bases of the oligonucleotides hold the same.

**Table S6. Sequences of the self-complementary RNA duplexes used in crystallographic studies.**

| Name | Source | Type | Sequence (5′→ 3′) [1] |
|---|---|---|---|
| UD-1 | In-house | RNA | AGA G<u>D</u>A GAU CU<u>U</u> CUC U |
| CD-1 | In-house | RNA | AGA G<u>D</u>A GAU CU<u>C</u> CUC U |
| UD-1 | In-house | RNA | AGA GAA G<u>DU</u> CUU CUC U |
| CD-2 | In-house | RNA | AGA GAA G<u>DC</u> CUU CUC U |

[1] Underlined bases indicate the probed base pair interactions.

**Table S7. Optimized Conditions for Crystallization**

| Sequence | Optimized crystallization conditions |
|---|---|
| UD1 | 80 mM Strontium chloride hexahydrate, 20 mM Magnesium chloride hexahydrate, 40 mM Sodium cacodylate trihydrate pH 7.0, 20% v/v (+/-)-2-Methyl-2,4-pentanediol, 12 mM Spermine. |
| UD2 | 40 mM Lithium chloride, 20 mM Magnesium chloride hexahydrate, 40 mM Sodium cacodylate trihydrate pH 5.5, 30% v/v (+/-)-2-Methyl-2,4-pentanediol, 2 mM Hexammine cobalt(III) chloride. |
| CD1 | 10 mM Magnesium chloride hexahydrate, 50 mM HEPES sodium pH 7.0, 4.0 M Lithium chloride. |
| CD2 | 400 mM Sodium chloride, 120 mM Calcium chloride, 27% v/v (+/-)-2-Methyl-2,4-pentanediol, 20 mM MES pH 5.8. |

**Table S8. Data Collection Statistics**

| Item | UD1 | UD2 | CD1 | CD2 |
|---|---|---|---|---|
| PDB code | 8TDY | 8TDZ | 8TE0 | 8TE2 |
| Beamline | 2.0.1 | 2.0.1 | 2.0.1 | 2.0.1 |
| Wavelength (Å) | 1.038413 | 1.038413 | 1.033216 | 1.038413 |
| Space group | R32 | R32 | R32 | R32 |
| Unit cell parameters (Å, °) | 41.15, 41.15, 125.05, 90, 90, 120 | 41.87, 41.87, 124.30, 90, 90, 120 | 42.50, 42.50, 129.98, 90, 90, 120 | 41.56, 41.56, 125.38, 90, 90, 120 |
| Resolution range (Å) | 50 – 1.65 (1.71 – 1.65) | 50 – 1.64 (1.67 – 1.64) | 50 – 1.42 (1.44 – 1.42) | 41.79 – 1.63 (1.66 – 1.63) |
| Unique reflections | 5058 (483) | 5403 (250) | 8586 (267) | 5477 (285) |
| Completeness (%) | 99.6 (99.2) | 99.6 (100) | 96.2 (64.5) | 99.7 (97.1) |
| Redundancy | 8.5 (6.1) | 8.7 (5.0) | 7.1 (2.0) | 8.3 (6.9) |
| $R_{merge}$ (%) | 9.7 (52.0) | 5.3 (18.4) | 6.5 (23.1) | 9.3 (36.2) |
| $\langle I/\sigma(I) \rangle$ | 17.0 (4.2) | 35.5 (8.4) | 30.0 (2.8) | 10.0 (2.4) |

**Table S9. Structure refinement statistics.**

| Item | UD1 | UD2 | CD1 | CD2 |
|---|---|---|---|---|
| PDB code | 8TDY | 8TDZ | 8TE0 | 8TE2 |
| Resolution range (Å) | 41.68 – 1.66 | 41.43 – 1.64 | 35.41 – 1.42 | 31.22 - 1.63 |
| Number of reflections | 4946 | 4953 | 7781 | 5344 |
| $R_{work}$ (%) | 19.6 | 18.8 | 21.6 | 21.0 |
| $R_{free}$ (%) | 22.4 | 22.2 | 23.8 | 25.2 |
| Bond length R.M.S. (Å) | 0.005 | 0.005 | 0.013 | 0.013 |
| Bond angle R.M.S. (°) | 0.95 | 1.00 | 2.37 | 2.10 |
| Average B-factors (Å$^2$) | 13.06 | 12.30 | 11.56 | 22.94 |

**Table S10. Sequences of the template and complementary oligonucleotides used in the deep sequencing experiments.**

| Name | Source | Type | Sequence (5′→ 3′) [1] |
|---|---|---|---|
| 6N | In-house | RNA/DNA | GUUCAGAGUUCUACAGUCCGACGAUCdT(-NPOM)CdT(NPOM)ANNNNNNGCAUGCGACUAAACGUCGCAUGC |
| 6D | In-house | RNA/DNA | GUUCDGDGUUCUDCDGUCCGDCGDUCdT(-NPOM)CdT(NPOM)DNNNNNNGCDUGCGDCUDDDCGUCGCDUGC |
| 5′ Handle Block | IDT | RNA | GUCGGACUGUAGAACUCUGAA-dideoxyC |
| RT Handle | IDT | DNA | App-AGATCGGAAGAGCACACGTCT-dideoxyC |
| RT Primer | IDT | DNA | AGACGTGTGCTCTTCCGATCT |
| PCR Primer 1 (SR Primer) | NEB | DNA | AATGATACGGCGACCACCGAGATCTACACGTTCAGAGTTCTACAG TCCG-s-A |
| PCR Primer 2 (Index Primer) | NEB | DNA | CAAGCAGAAGACGGCATACGAGAT(6-base index)GTGACTGGAGTTCAGACGTGTGCTCTTCCGATC-s-T |

**Table S11. Overall frequencies of total extended products in AUCG and DUCG systems, and their ratios.**

| Reaction conditions | AUCG | DUCG | DUCG/AUCG |
|---|---|---|---|
| 10 mM $MgCl_2$, 10 mM N*N | 0.15 | 0.21 | 1.38 |
| 10 mM $MgCl_2$, 20 mM N*N | 0.21 | 0.18 | 0.86 |
| 100 mM $MgCl_2$, 20 mM N*N | 0.31 | 0.42 | 1.36 |
